# Structural insights into the agonist selectivity and structure-based engineering of the adenosine A3 receptor

**DOI:** 10.1101/2024.03.23.586386

**Authors:** Hidetaka S. Oshima, Akiko Ogawa, Fumiya K. Sano, Hiroaki Akasaka, Tomoyoshi Kawakami, Hiroyuki H. Okaomto, Aika Iwama, Chisae Nagiri, Fan-Yan Wei, Wataru Shihoya, Osamu Nureki

## Abstract

Adenosine receptors, expressed across various tissues, play pivotal roles in physiological processes and are implicated in diverse diseases, including neurological disorders and inflammation, highlighting the therapeutic potential of receptor-selective agents. The Adenosine A3 receptor (A_3_R), the last identified adenosine receptor, is also activated by breakdown products of post-transcriptionally modified tRNA and exhibits dual roles in neuron, heart, and immune cells, and is often overexpressed in tumors, making it a target for anticancer therapy. Despite extensive studies on the other adenosine receptors, the structure and activation mechanism of A_3_R, especially by selective agonists like *N*^6^-methyladenosine (m^6^A) and namodenoson, remained elusive. Here, we identified *N*^6^-isopentenyl adenosine (i^6^A), a novel A_3_R-selective ligand, via comprehensive modified adenosine library screening. Cryo-EM analyses of A_3_R-G_i_ signaling complexes with two nonselective and three selective agonists revealed the structural basis for A_3_R activation. We further conducted structure-guided engineering of m^6^A-insensitive A_3_R, which would greatly facilitate future discoveries of the physiological functions of the selective activation of A_3_R by modified adenosines. Our results clarify the selective activation of adenosine receptors, providing the basis for future drug discovery.

## Introduction

Adenosine functions as an extracellular signaling molecule by activating adenosine receptors, besides its fundamental role as a building block of RNA. Adenosine receptors belong to the class A G-protein coupled receptors (GPCRs), and are expressed in various types of tissues(Fredholm et al., 2001). Adenosine-mediated receptor activation engages in various physiological responses, including immunity, sensory conception, learning, and memory. Furthermore, the involvement of adenosine receptors has been reported in various diseases, including mental disorders and inflammation(Effendi et al., 2020). In those diseases, the roles of each receptor subtype are diverse; for example, inhibition of the A_2A_R has been effective in treating Parkinson’s disease, while the activation of A_1_R has helped prevent inflammation-mediated organ injury in sepsis(Cieślak et al., 2008; Gallos et al., 2005). Therefore, the development of selective agents targeting adenosine receptors has garnered keen attention.

Of the four adenosine receptors, A_3_R is the last to be identified(Zhou et al., 1992). A_3_R is widely distributed throughout the body, including in neuron, heart, and immune cells, and is most highly expressed in the lungs and liver. A_3_R activation reportedly plays dual roles, offering both neuroprotection and neurodegeneration, cardioprotection and cardiotoxicity, and anti-inflammatory and proinflammatory effects(Gessi et al., 2008). Thus, the role of A_3_R in disease remains an area of intense research. Using extracellular modified nucleosides as ligands, we have recently identified m^6^A as a selective and potent ligand for A_3_R, with potency 10-fold stronger than that of adenosine(Ogawa et al., 2021). m^6^A is one of the RNA modifications which exists in most RNA species including mRNA, rRNA and tRNA, and is released as modified nucleosides after the catabolic process of RNA degradation. Elevation of extracellular m^6^A concentrations effectively induced inflammation and type I allergy through A_3_R. Moreover, it is particularly noteworthy that A_3_R is often overexpressed in tumors(Madi et al., 2004). The activation of A_3_R in tumor cells is associated with anticancer effects, and A_3_R-selective agonists such as namodenoson and piclidenoson are currently in clinical trials and showing promising results(Fishman, 2022).

Although numerous structures of the other adenosine receptors have been reported, the structure of the A_3_R has remained enigmatic(Cai et al., 2022; Carpenter et al., 2016; Chen et al., 2022; Cheng et al., 2017; Doré et al., 2011; Draper-Joyce et al., 2021, 2018; Glukhova et al., 2017; Jaakola et al., 2008; Xu et al., 2011). Consequently, the specific mechanisms through which nonselective and selective agonists activate A_3_R are still unclear. Moreover, details about the interactions between A_3_R and modified adenosines, like m^6^A, have yet to be clarified. Here, we discovered a new A_3_R-selective ligand, i^6^A, through screening using a comprehensive modified adenosine library. Subsequent cryo-EM structural analyses elucidated the structures of the A_3_R-G_i_ complex bound to two nonselective and three selective agonists, revealing the structural basis of the agonist-bound activation of A_3_R.

## Results

### Comprehensive screening of modified nucleosides against human A_3_R

Using a limited set of modified nucleosides, we previously showed that m^6^A selectively activates A_3_R(Ogawa et al., 2021). To fully understand the functional roles of RNA-derived modified nucleosides with A_3_R, we tested as many as 42 species of modified nucleosides with A_3_R as well as other adenosine receptors in TGFα shedding assays(Inoue et al., 2012) (Figures 1A and 1B; Table S1). Consistent with the previous study, m^6^A showed the greatest potency only for A_3_R compared with other modified nucleosides and adenosine (calculated intrinsic activity (RAi) values of m^6^A relative to adenosine = 13 ± 3.6(Ogawa et al., 2021)) (Figure 1C). In addition to m^6^A, we found that *N*^6^-isopentenyladenosine (i^6^A), a tRNA-specific RNA modification, selectively activates A_3_R (RAi = 0.59 ± 0.046) (Figure 1D). Moreover, *N*^6^, *N*^6^-dimethyladenosine (m^6,6^A), which is the dimethyl variant of m^6^A exclusively found in 18S rRNA, showed very weak activation of A_3_R (RAi = 0.069 ± 0.0057) (Figure 1E). Interestingly, *N*^6^-threonylcarbamoyladenosine (t^6^A), another tRNA-specific RNA modification at the *N*^6^ position, did not show any potency on A_3_R (Figure 1F). These results suggest that the chemical properties of the modification at the *N*^6^ position profoundly impact the receptor activation potency.

**Figure 1.**
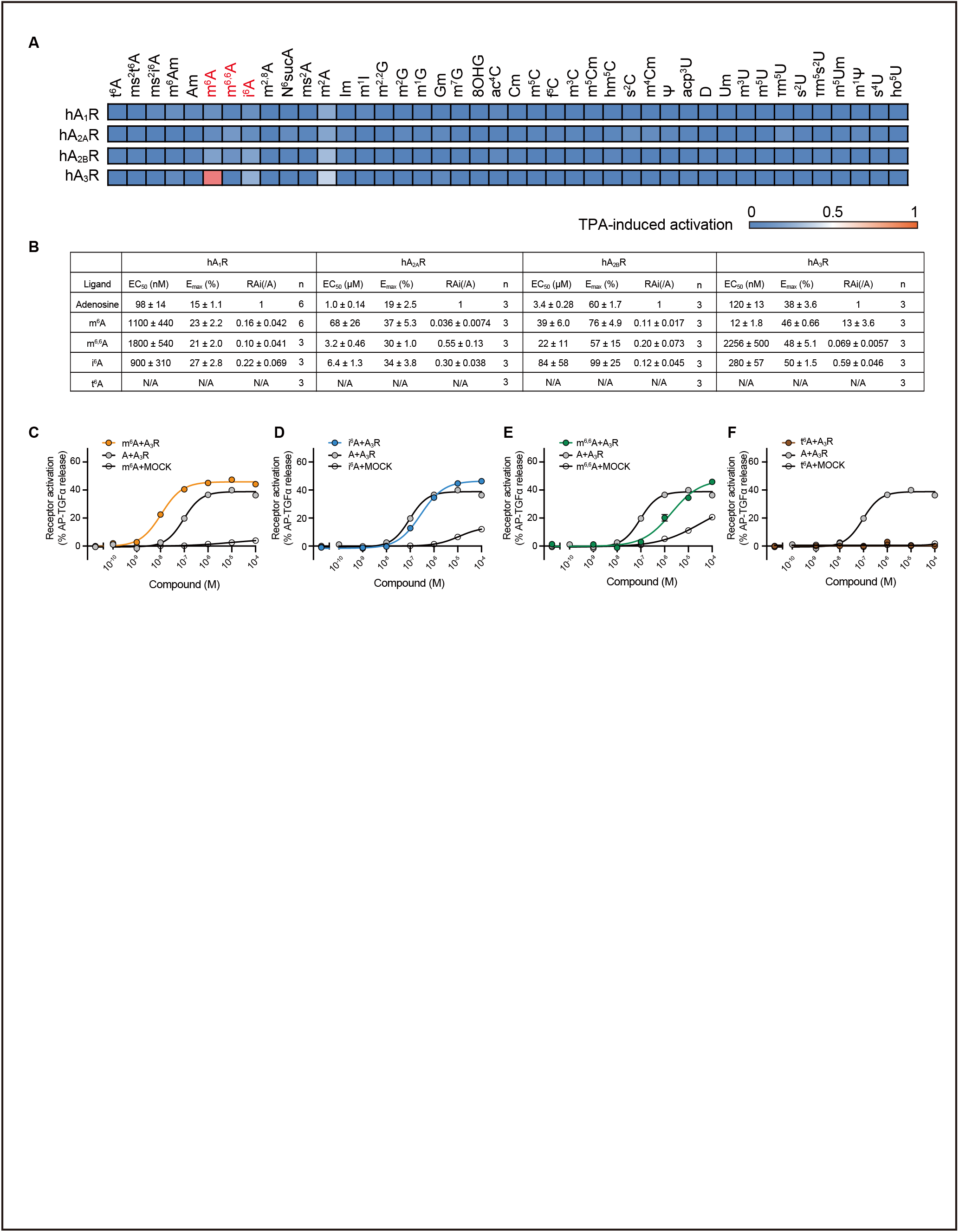
Comprehensive screening of modified nucleosides against human A_3_R. (A) Heatmaps showing the activation of adenosine receptor subtypes by modified nucleosides, as measured by the TGFα-shedding assay. The color scale represents % receptor activation compared to TPA (12-O-tetradecanoylphorbol 13-acetate)-mediated receptor activation, which induces the maximum TGFα-shedding response independently of the receptor, and the adenosine receptor-dependence of signals were calculated by subtracting the response in mock-transfected cells from the response in adenosine receptor-expressing cells. The tested compounds’ concentrations were 100 nM for hA_1_R and hA_3_R, 1 µM for hA_2A_R, and 5 µM for hA_2B_R (n = 3) (B) EC_50_ and E_max_ values of adenosine, *N*^6^-methyladenosine (m^6^A), *N*^6^, *N*^6^-dimethyladenosine (m^6,6^A), *N*^6^-isopentenyladenosine (i^6^A), and *N*^6^-threonylcarbamoyladenosine (t^6^A) for adenosine receptor subtypes, as determined by the TGFα-shedding assay. Data are shown as means ± SEM, n = 3 unless otherwise noted. (C-F) Comparison of TGFα-shedding response curves for hA_3_R between modified nucleosides (C: m^6^A, D: m^6,6^A, E: i^6^A, and F: t^6^A) and adenosine. Response curves for mock-transfected cells are shown in the same graph (Data are shown as means ± SEM, n = 3).

It should be noted that we observed that 1-methyladenosine (m^1^A), which is one of the major and abundant RNA modifications, can also induce A^3^R activation. However, since synthetic m^1^A contains a significant amount of m^6^A as a by-product of chemical synthesis(Liu et al., 2022), we did not include m^1^A in the results. Additionally, 2-methyladenosine (m^2^A) showed weak and non-selective potency towards all four adenosine receptors, although m^2^A is only found in *Escherichia coli* and plants, unlike m^6^A, i^6^A, m^6,6^A, and t^6^A, which are found in humans.

### Overall structure of the A_3_R-G_i_ complex

For the structural analysis, we focused not only on the modified adenosine analogues obtained through the screening but also on adenosine, 5’-N-ethylcarboxamidoadenosine (NECA), and namodenoson. NECA is an adenosine analogue that acts as a potent agonist for adenosine receptors, and several structures of adenosine receptors bound to NECA have been reported(Carpenter et al., 2016; Chen et al., 2022; García-Nafría et al., 2018; Lebon et al., 2011). Namodenoson is an A_3_R-selective agonist with potential efficacy in cancer treatment(Fishman, 2022). We used these two nonselective (adenosine and NECA) and three (m^6^A, i^6^A and namodenoson) selective agonists for our structural analysis, but excluded m^6,6^A due to its low potency and weak selectivity (Figure 2A).

**Figure 2.**
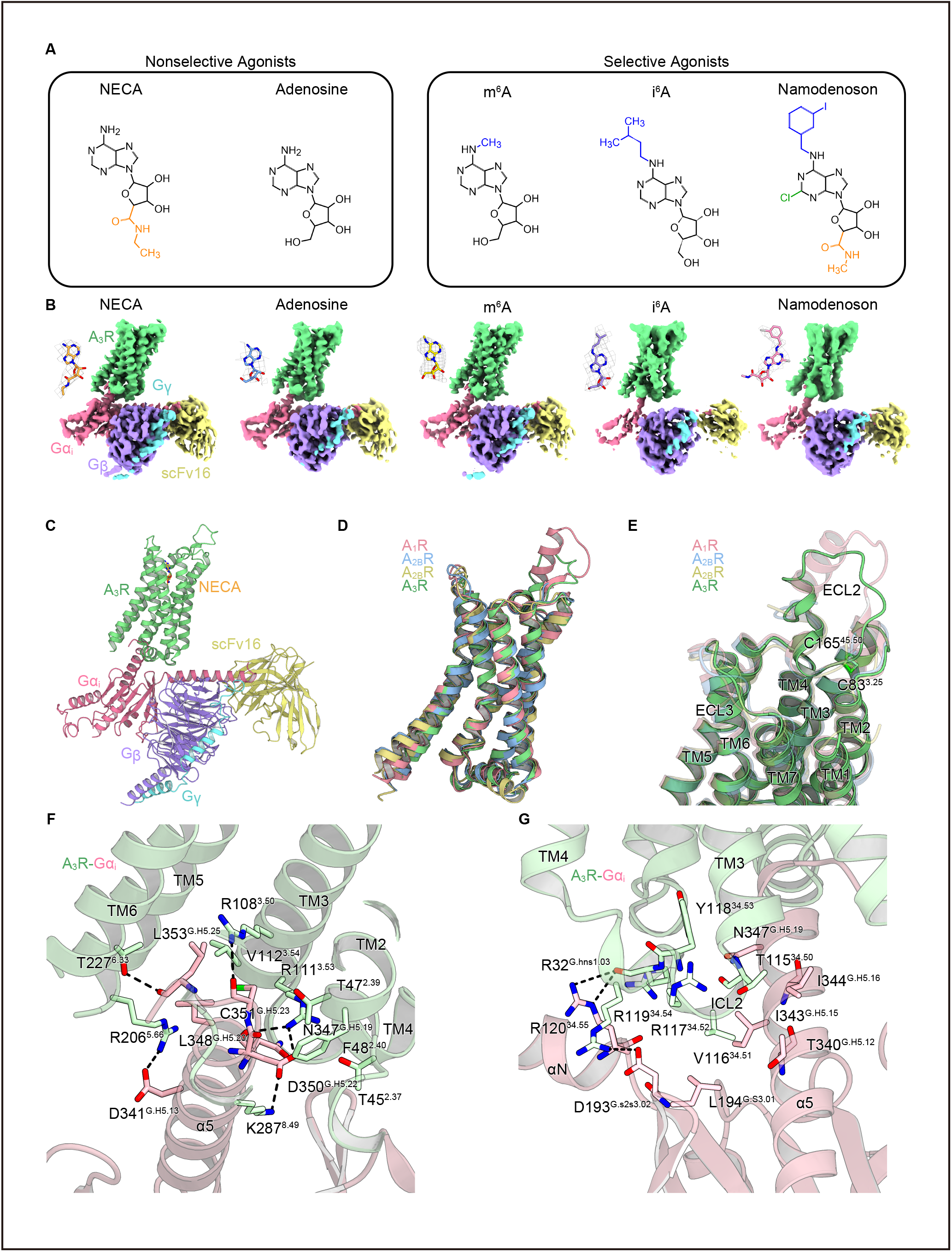
Overall structure of the A_3_R-G_i_ complex. (A) Structures of the agonists used for structural analyses. (B) Cryo-EM maps of the A_3_R-G_i_ complex bound to NECA, adenosine, m^6^A, i^6^A, and namodenoson. Densities of each agonist are shown in the top-left corner of each map. (C) Overall structure of the A_3_R-G_i_ complex bound to NECA. (D) Superimposition of A_1_R (PDB 6D9H), A_2A_R (PDB 6GDG), A_2B_R (PDB 8HDP), and A_3_R. (E) Close-up view of ECLs of adenosine receptors. ECL2 and ECL3 show remarkable differences among the adenosine receptors. The disulfide bond between ECL2 and TM3 of A_3_R is labeled. (F-G) Receptor-G-protein interactions around α5 (F) and ICL2 (G). The residues involved in the interactions are represented by stick models. Black dashed lines indicate hydrogen bonds.

Initially, we attempted the structural analysis of a human A_3_R, but it showed poor monodispersity in fluorescence size exclusion chromatography (FSEC)(Hattori et al., 2012). Thus, we evaluated vertebrate A_3_R homologues by FSEC. Consequently, we identified sheep A_3_R as the most suitable candidate for structural analysis (Figure S1A). The transmembrane regions of sheep and human A_3_R exhibit high sequence homology of 90%(Pándy-Szekeres et al., 2018; Robert and Gouet, 2014; The UniProt Consortium, 2023) (Figure S1B). To examine the ligand binding profile of sheep A_3_R, we evaluated the agonist efficacies of the human and sheep A_3_Rs by a TGFα shedding assay(Inoue et al., 2012), and found that they are quite similar (Figure S1C; Table S2). Moreover, the mRNA distribution of sheep A_3_R is similar to that in humans, with expression in the lungs and pineal glands(Gao et al., 2023). Considering these factors, we concluded that the functional properties of the human and sheep A_3_Rs do not differ significantly, and proceeded with the structural analysis of the sheep A_3_R-G_i_ complexes.

For sample preparation, we adopted the previously reported Fusion-G system, which combines the NanoBiT tethering system and the fusion of the G_α_ and G_γ_ subunits as a single polypeptide(Duan et al., 2020; Kim et al., 2020; Nureki et al., 2022; Sano et al., 2023). We co-expressed sheep A_3_R and the fusion-G_i_ trimer in HEK293 cells, solubilized the complex with detergent, and then purified it using FLAG affinity chromatography. The complex was stabilized with a single-chain variable fragment (scFv16), isolated via gel filtration chromatography (Figure S1D), and subsequently used for cryo-EM analysis (Figures S2A-S2E). Finally, we obtained the cryo-EM maps of the A_3_R complex bound to the two nonselective and three selective agonists at overall resolutions of 2.86 Å (NECA), 3.27 Å (adenosine), 3.40 Å (m^6^A), 3.66 Å (i^6^A) and 3.62 Å (namodenoson) (Figure 2B). Notably, the NECA-bound complex exhibited the highest resolution, and thus this structure is used in the following discussion of the overall structure (Figure 2C).

Comparing the overall structure of A_3_R with the cryo-EM structures of other adenosine receptors, A_3_R overlaps with A_1_R, A_2A_R, and A_2B_R with root-mean-square deviations (R.M.S.Ds) of 0.82, 0.96, and 0.90 Å, respectively, indicating considerable similarity with A_1_R(Cai et al., 2022; Draper-Joyce et al., 2018; García-Nafría et al., 2018) (Figure 2D). While the overall structures of the adenosine receptors superimpose well, extracellular loops (ECLs) 2 and 3 exhibit secondary structure-level differences (Figure 2E). ECL2 of the adenosine receptors has varied sequences across the family, and that of A_3_R is the shortest among them (Figure S3). From the N- to C-terminal side, ECL2 of the adenosine receptors comprises a short helical region, a disordered region, a sheet-like region forming a disulfide bond with the transmembrane helix (TM) 3, and a comparatively conserved region that contributes to the ligand pocket. ECL2 of A_3_R forms a relatively shorter helix compared to A_1_R on the N-terminal side (Figure 2E). We were able to model the subsequent disordered region, which is also short and distant from the other regions, implying little relevance to the receptor function. In the following sheet-like region, C83^3.25^ and C165^45.50^ form a disulfide bond (superscripts indicate Ballesteros–Weinstein numbers(Ballesteros and Weinstein, 1995)), as in the other adenosine receptors. Furthermore, ECL3 of A_3_R is the shortest among the adenosine receptors, lacking the N-terminal helical structure observed in the other receptors and instead extending straight towards TM7.

Next, we inspected the interaction between A_3_R and G_i_. There are hallmark regions of the interactions between class A G_i_-coupling receptors, including A_3_R, and G_i_: ICL2 of a receptor as well as α5 of G_i_(Akasaka et al., 2022; Hua et al., 2020; Izume et al., 2024; Kato et al., 2019; Okamoto et al., 2021; Oshima et al., 2024; Sano et al., 2023; Yuan et al., 2021; Zhuang et al., 2022). In detail, around α5 of G_i_, several residues form hydrogen bonds, represented by the interaction between R108^3.50^ and C351^H5.23^ (superscript indicates the common Gα numbering [CGN] system(Flock et al., 2015)) (Figure 2F). In the ICL2 of A_3_R, V116^34.51^ penetrates into the hydrophobic cavity of G_i_, which is common in class A GPCRs(Akasaka et al., 2022; Hua et al., 2020; Iwama et al., 2023; Izume et al., 2024; Kato et al., 2019; Nagiri et al., 2021; Okamoto et al., 2021; Oshima et al., 2024; Rasmussen et al., 2011; Sano et al., 2023; Yuan et al., 2021; Zhuang et al., 2022) (Figure 2G). Furthermore, extensive ionic interactions are formed across ICL2. These extended interactions more closely resemble that between A_2B_R and G_s_, rather than that between A_1_R and G_i_(Cai et al., 2022; Chen et al., 2022; Draper-Joyce et al., 2021, 2018; García-Nafría et al., 2018; Zhang et al., 2022) (Figures S4A-S4D). Moreover, A_3_R has T115^34.50^ at the C-terminus of TM3, while most class A GPCRs including the other adenosine receptors have P^34.50^ at this position. This results in a slightly elongated TM3, possibly enabling the extensive interactions across ICL2.

### Binding modes of adenosine and NECA

The orthosteric ligand pocket of A_3_R consists of the extracellular halves of TM3, TM5, TM6, TM7, and ECL2, as in the other adenosine receptors(Cai et al., 2022; Chen et al., 2022; Draper-Joyce et al., 2021, 2018; García-Nafría et al., 2018; Zhang et al., 2022) (Figure 3A). The binding modes of adenosine and NECA are remarkably similar in terms of the adenosine moiety (Figures 3B and 3C). In detail, the adenine moiety forms hydrogen bonds with N249^6.55^ and pi-stacking interactions with F167^45.52^, while the ribose moiety forms hydrogen bonds with H271^7.43^, and there are extensive hydrophobic interactions surrounding the entire ligand. Moreover, the modification at the 5’ position of NECA forms an extra hydrogen bond with T94^3.36^, and the following carbon chain engages in hydrophobic contacts with the W242^6.48^, which is known as the toggle switch motif essential for class A GPCR activation(Zhou et al., 2019), and residues in its vicinity (Figure 3C). These additional interactions may stabilize NECA more effectively than adenosine, enabling the activation of A_3_R at lower concentrations as in other adenosine receptors.

**Figure 3.**
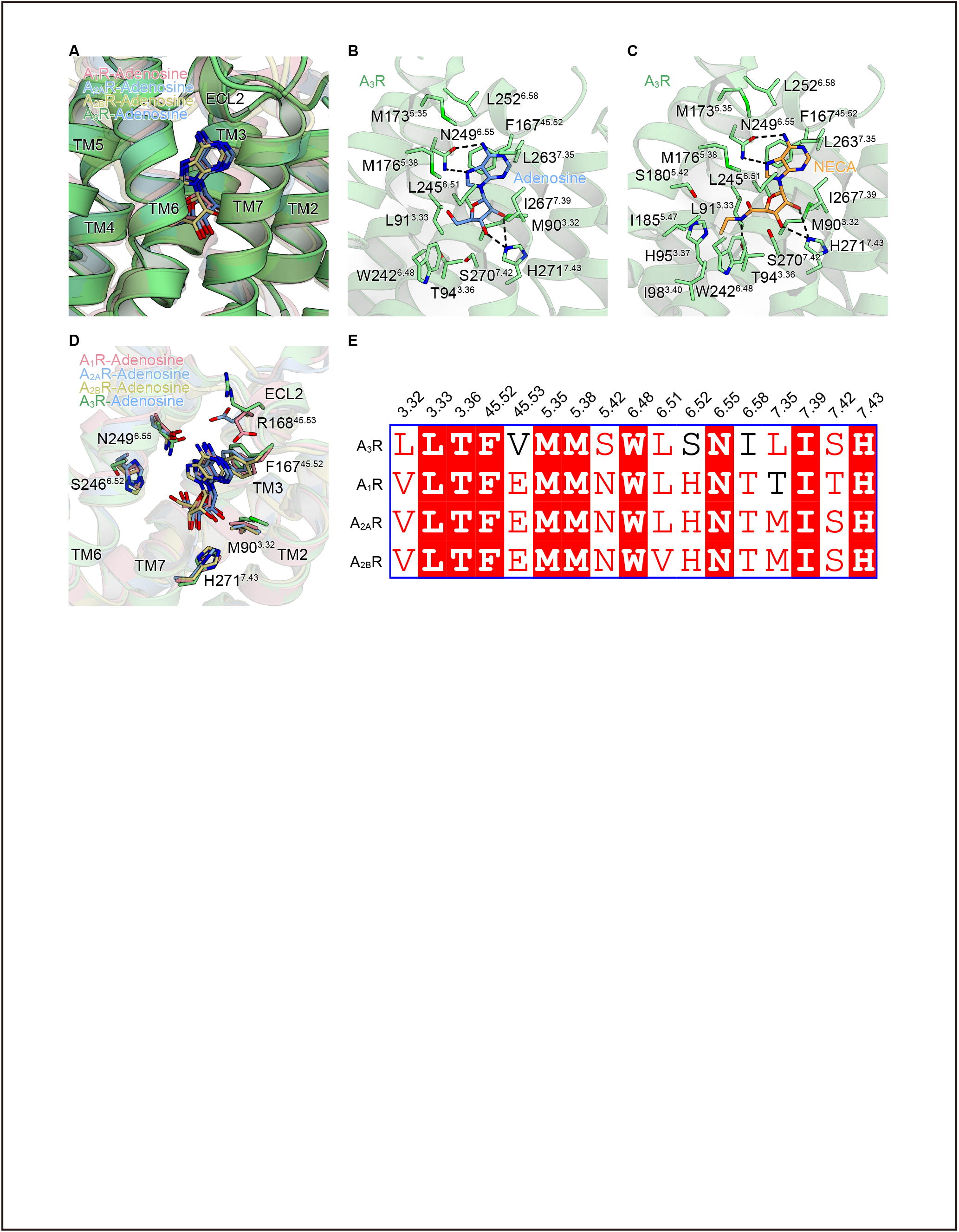
Binding modes of adenosine and NECA. (A) Superimposition of A_1_R (PDB 6D9H), A_2A_R (PDB 2YDO), A_2B_R (PDB 8HDP), and NECA-bound A_3_R. The adenosine moieties of the ligands are accommodated in a similar position. (B-C) Binding modes of adenosine (B) and NECA (C). Black dashed lines indicate hydrogen bonds. (D) Comparison of the binding modes of ligands to each adenosine receptor. Representative residues involved in the ligand-receptor interaction are represented by stick models. Residues of A_3_R are labeled. Key interactions are conserved among adenosine receptors. (E) Alignment of residues comprising the orthosteric pockets of adenosine receptors.

Most residues in the orthosteric pocket are conserved, and the binding sites and binding modes of the adenosine moiety are quite similar among the adenosine receptors, including key interactions such as hydrogen bonds with N^6.55^ and H^7.43^, and pi-stacking interactions with F^45.52^ (Ref. 11, 14, 16, 20, 30, 31) (Figures 3D and 3E). Nevertheless, there are subtle differences between A_3_R and other adenosine receptors. The residue at 3.32 is leucine in human A_3_R, and methionine in sheep A_3_R. Both are longer than the valine of the other adenosine receptors, probably resulting in more intimate contact with a ligand in A_3_R. At position 6.52, while the other adenosine receptors feature histidine, which electrostatically interacts with the 5’ hydroxyl group of the adenosine moiety, A_3_R possesses serine, which does not participate in the ligand-receptor interaction. At position 45.53 in ECL2, the other adenosine receptors have a glutamate that forms a hydrogen bonding interaction with the adenine moiety. Notably, a mutation at this position in A_2A_R notably led to a significant decrease in its activity(Guo et al., 2016). By contrast, human and sheep A_3_R have valine and arginine at this position, respectively, and V169^45.53^E mutation was reported to greatly reduce the potency of adenosine(Ogawa et al., 2021). Despite the absence of glutamate at this position, adenosine showed higher potency for A_3_R than for A_2A_R or A_2B_R, indicating that A_3_R has unique features not found in other adenosine receptors.

### Binding modes of m^6^A and i^6^A

m^6^A binds to A_3_R in the same position as adenosine, exhibiting the common interactions such as hydrogen bonding with N249^6.55^ and H271^7.43^, and pi-stacking interactions with F167^45.52^ (Figures 4A and 4B). Moreover, the methyl group at the *N*^6^ position forms close hydrophobic interactions with the aliphatic side chains of R168^45.53^, M173^5.35^, and L252^6.58^ (Figure 4C). These residues create a tightly packed hydrophobic pocket along with the nearby L263^7.35^, although the latter is not in direct interaction distance with the methyl group. In contrast to A_3_R, the residues at these positions are hydrophilic in the other adenosine receptors. Previously, mutations of these A_3_R residues to the corresponding residues in the other adenosine receptors significantly decreased the potency of m^6^A(Ogawa et al., 2021), indicating that these hydrophobic residues exactly enable m^6^A to selectively activate A_3_R. Furthermore, A_3_R exhibits much tighter packing in this pocket than the other adenosine receptors (Figure 4D). This narrow hydrophobic pocket fits very well with the methyl group of m^6^A, probably contributing to the high and selective potency of m^6^A. Moreover, this tight packing on the extracellular side of the ligand pocket may explain the comparably higher potency of adenosine for A_3_R.

**Figure 4.**
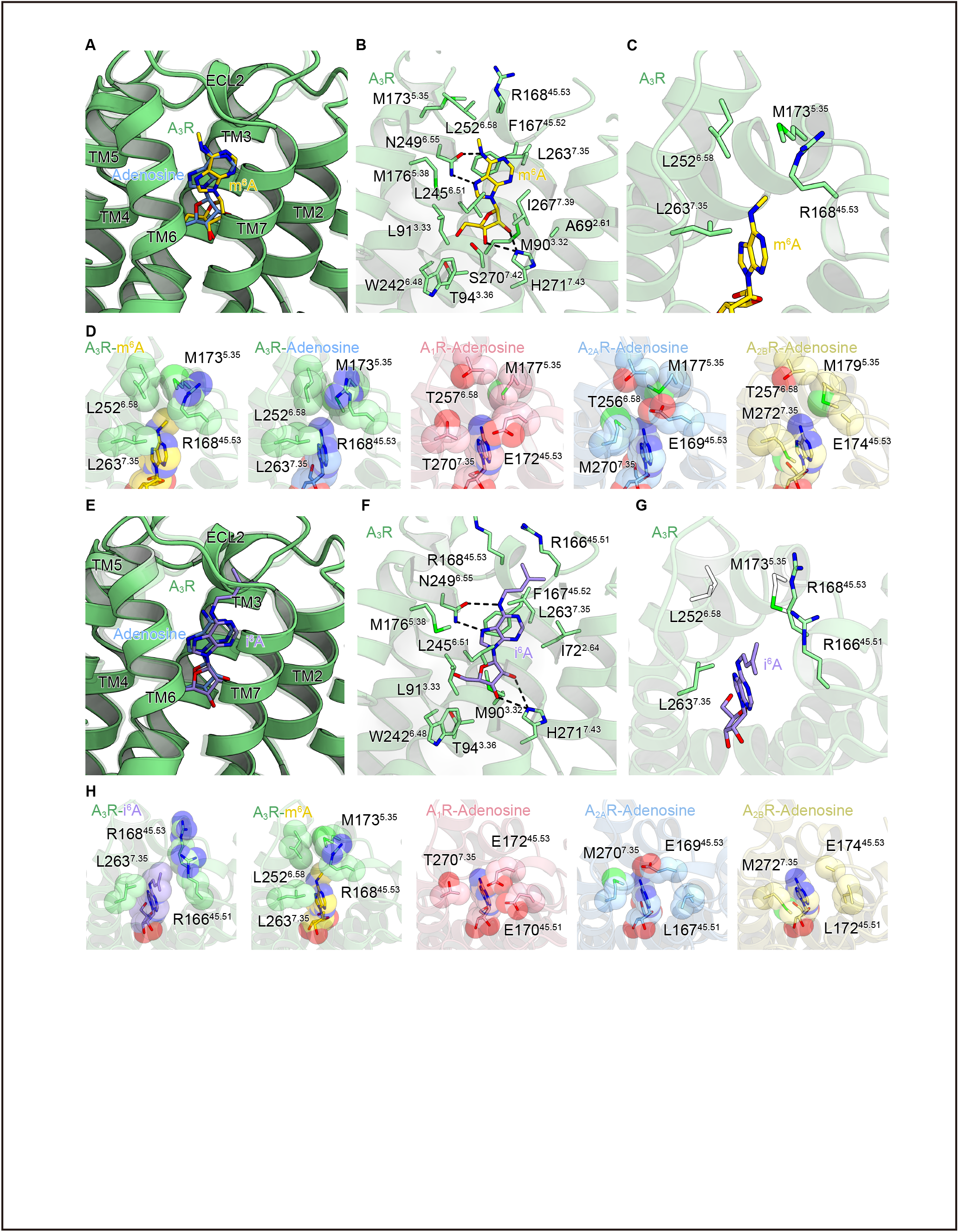
Binding modes of m^6^A and i^6^A. (A) Superimposition of adenosine-bound and m^6^A-bound A_3_R. The adenosine-bound A_3_R is represented by a translucent ribbon model. (B) Binding mode of m^6^A. Black dashed lines indicate hydrogen bonds. (C) Close-up view of the hydrophobic pocket of the m^6^A-bound model. (D) CPK models of the hydrophobic pocket of A_3_R and those of A_1_R, A_2A_R, A_2B_R, and adenosine-bound A_3_R (PDB 6D9H, 2YDO, 8HDP, and this study, respectively). The methyl group of m^6^A is especially tightly packed. (E) Superimposition of adenosine- and i^6^A-bound A_3_R. The adenosine-bound receptor is represented by a translucent ribbon model. (F) Binding mode of i^6^A. Black dashed lines indicate hydrogen bonds. (G) Close-up view of the hydrophobic pocket of the i^6^A-bound structure. The isopentyl group of i^6^A diverges from the hydrophobic pocket and forms hydrophobic interactions with R166^45.51^, R168^45.53^ and L263^7.35^. M173^5.35^ and L252^6.58^ are not close enough to interact with i^6^A (colored white). (H) CPK models of the residues of A_3_R interacting with the 2-isopentyl group of i^6^A and those of A_1_R, A_2A_R, A_2B_R, and m^6^A-bound A_3_R (PDB 6D9H, 2YDO, 8HDP, and this study, respectively).

The adenosine moiety of i^6^A is also accommodated in a comparable position to adenosine, and forms similar interactions to other agonists (Figures 4E and 4F). However, in contrast to m^6^A, the isopentyl group at the *N*^6^ position is diverged from the hydrophobic pocket and oriented toward R166^45.51^, forming hydrophobic interactions with the aliphatic side chains of R166^45.51^, R168^45.53^, and L263^7.35^, but not with M173^5.35^ and L252^6.58^ (Figure 4G). In the other adenosine receptors, E^45.53^ forms a hydrogen bond with the 6-amino group of the adenine moiety, making it difficult for modifications at the *N*^6^ position to orient towards 45.51 and extend extracellularly (Figure 4H). This could explain why i^6^A is selective for A_3_R. Moreover, the interaction between i^6^A and the receptor is looser compared to that of m^6^A, which may explain the significantly lower potency of i^6^A towards A_3_R than that of m^6^A.

### Structure-guided engineering of m^6^A-insensitive A_3_R

We previously reported that the selective activation of A_3_R by m^6^A induces allergies and inflammation *in vivo*(Ogawa et al., 2021). However, the biological phenomena involving A_3_R also include neurotransmission, cell proliferation, apoptosis, and so forth(Gessi et al., 2008). To thoroughly investigate the role of the selective activation of A_3_R by m^6^A, to develop an engineered A_3_R insensitive to m^6^A but still sensitive to adenosine would be quite helpful, since it would be able to knock out the m^6^A-mediated pathway while maintaining the intact adenosine-mediated pathway. Thus, we generated comprehensive mutants of the residues forming the hydrophobic pocket that accommodates the methyl group of m^6^A; namely, V169^45.53^, M174^5.35^, I253^6.58^, and L264^7.35^ in human A_3_R (Figure 5A). We evaluated these mutants by monitoring the changes of the RAi values in the TGFα shedding assay(Inoue et al., 2012).

**Figure 5.**
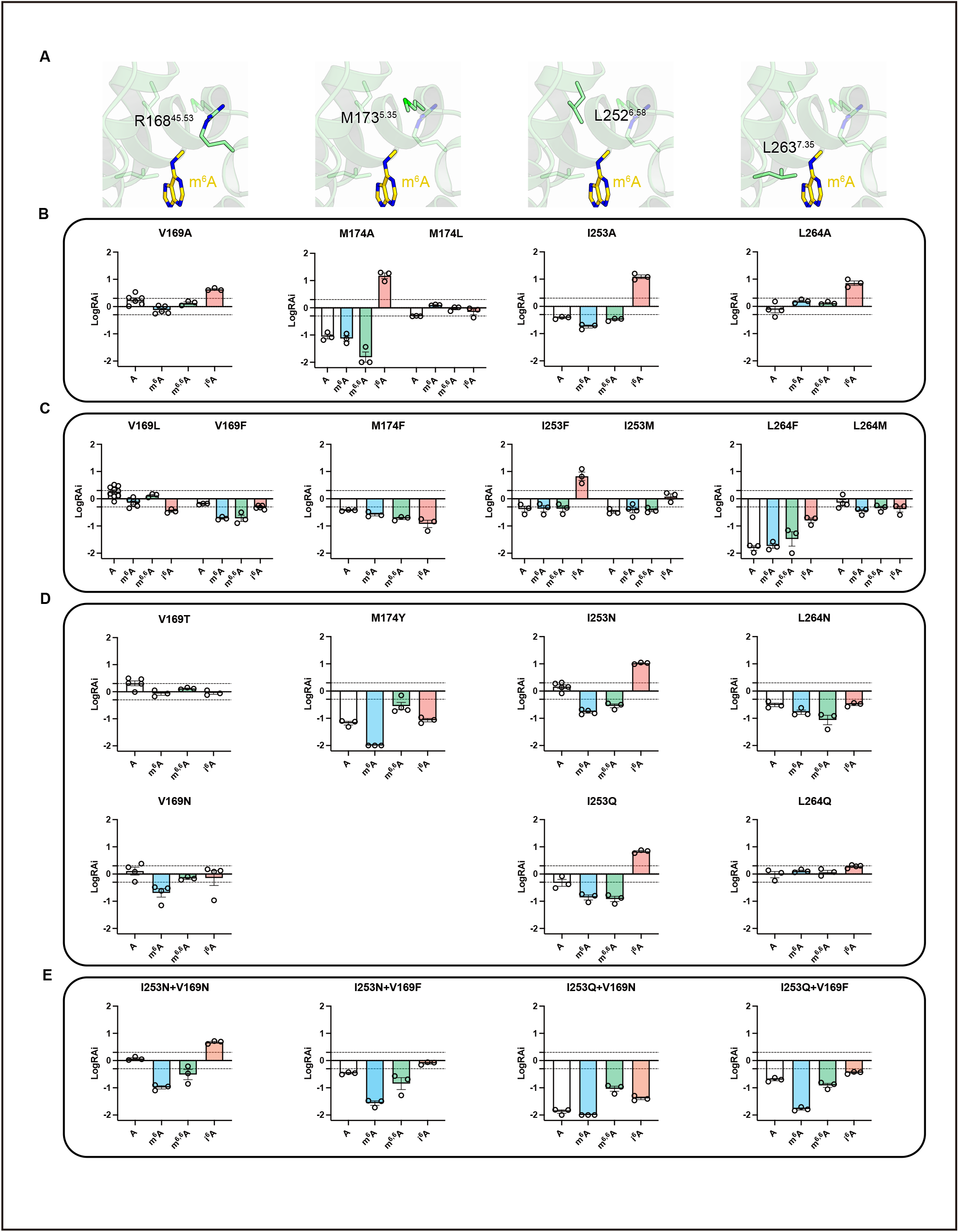
Structure-guided engineering of m^6^A-insensitive A_3_R. (A) Close-up views of the hydrophobic pocket of the m^6^A-bound A_3_R structure. Each column corresponds to mutations to the displayed residue. (B-E) Relative RAi values of adenosine, m^6^A, m^6,6^A, and i^6^A for human A_3_R mutants, as determined by the TGFα-shedding assay. RAi values are expressed as fold change of the values for the WT. Data are shown as means ± SEM (n = 3). Mutations that enlarge (B) and reduce (C) the size of the hydrophobic pocket. (D) Mutations that replace the original residue to a hydrophilic residue. (E) Designed mutants for selectively reducing the potency of m^6^A.

We initially examined mutations that change the side chain bulkiness of these residues (Figures 5B and 5C). We speculated that reducing the bulkiness of the residues would reduce the potency of m^6^A, since the tight hydrophobic interaction between A_3_R and the methyl group of m^6^A would be lost. Surprisingly, the potency of m^6^A was only reduced in the M174^5.35^A and I253^6.58^A mutants (Figure 5B). Since adenosine was also less potent in these mutants, this reduced potency of m^6^A may stem from the disruption of the packing in the hydrophobic pocket, rather than the loss of the hydrophobic interaction with the methyl group. By contrast, the potency of i^6^A generally increased among these mutants. Given that i^6^A has a bulkier modification at the *N*^6^ position compared to m^6^A, these mutants may create another space within the hydrophobic pocket that is favorable for i^6^A.

Next, we examined mutations that increase the side chain bulkiness of these residues. We speculated that a bulkier side chain would overlap with an *N*^6^ modification, reducing the potency of the modified adenosines. Consistently, these mutations generally reduced the potency of all ligands (Figure 5C). This is probably because the larger side chains occlude the pocket, in contrast to the enlargement of the pocket. However, some mutants exhibited remarkable behaviors. V169^45.53^L increased the potency of adenosine. This is probably because the extended carbon chain in the V169^45.53^L mutation occludes the space that accommodates the methyl group of m^6^A, compensating for the tight packing. Among these mutants, V169^45.53^F showed a clearer decrease in the potency of m^6^A compared to adenosine, and thus is a good candidate for an engineered A_3_R.

Subsequently, we changed residues in the hydrophobic pocket to hydrophilic residues (Figure 5D). We speculated that hydrophilic side chains would reduce the potency of the hydrophobically modified adenosines. We created two types of mutants: ones with unchanged side chain bulk and others with increased side chain bulk. As expected, these mutants generally exhibited decreased potency with m^6^A. However, the effects on the potency of adenosine varied depending on the mutation. Among these mutants, I253^6.58^N, I253^6.58^Q, and V169^45.53^N showed significantly larger drops in the potency of m^6^A than adenosine, highlighting their potential as components of functional mutants.

Based on these results, we combined the promising mutants identified in our experiments; namely, I253^6.58^N, I253^6.58^Q, V169^45.53^N, and V169^45.53^F (Figure 5E). Among the four combinations of these mutants, I253^6.58^N/V169^45.53^N and I253^6.58^N/V169^45.53^F especially caught our attention. While I253^6.58^N/V169^45.53^N significantly reduced the potency of m^6^A, it retained the potency of adenosine. I253^6.58^N/V169^45.53^F reduced the potency of m^6^A even more than I253^6.58^N/V169^45.53^N. Although I253^6.58^N/V169^45.53^F also slightly decreased the potency of adenosine, considering that the concentrations of adenosine and m^6^A are about 150 nM and 30 nM in human plasma(Ogawa et al., 2021), this mutant could have just the right affinity to almost completely shut off the m^6^A pathway under *in vivo* conditions, while minimally affecting the adenosine pathway. These two mutants would greatly facilitate future discoveries of the physiological functions of the selective activation of A_3_R by m^6^A.

### Binding mode of namodenoson

Namodenoson has an adenosine backbone with a 3-iodobenzyl group at the *N*^6^ position, a chloro group at the 2 position, and a methylamino group at the 5’ position (Figure 2A). The adenosine moiety of namodenoson is accommodated in the same position as the other agonists, by hydrogen bonding with N249^6.55^ and H271^7.43^, pi-stacking interactions with F167^45.52^, and extensive hydrophobic interactions (Figures 6A and 6B). Like the methyl group of m^6^A, the 3-iodobenzyl group at the *N*^6^ position interacts with the residues in the hydrophobic pocket. Moreover, it wedges into the hydrophobic pocket and exhibits more extensive hydrophobic interactions than m^6^A (Figure 6C). In the past structure-activity relationship (SAR) studies, the 3-iodobenzyl group at the *N*^6^ position strongly enhanced the A_3_R selectivity(Gallo-Rodriguez et al., 1994). As in m^6^A, the modification at the *N*^6^ position interacts extensively and closely with an A_3_R-specific hydrophobic pocket, likely endowing namodenoson with its strong and selective potency towards A_3_R.

**Figure 6.**
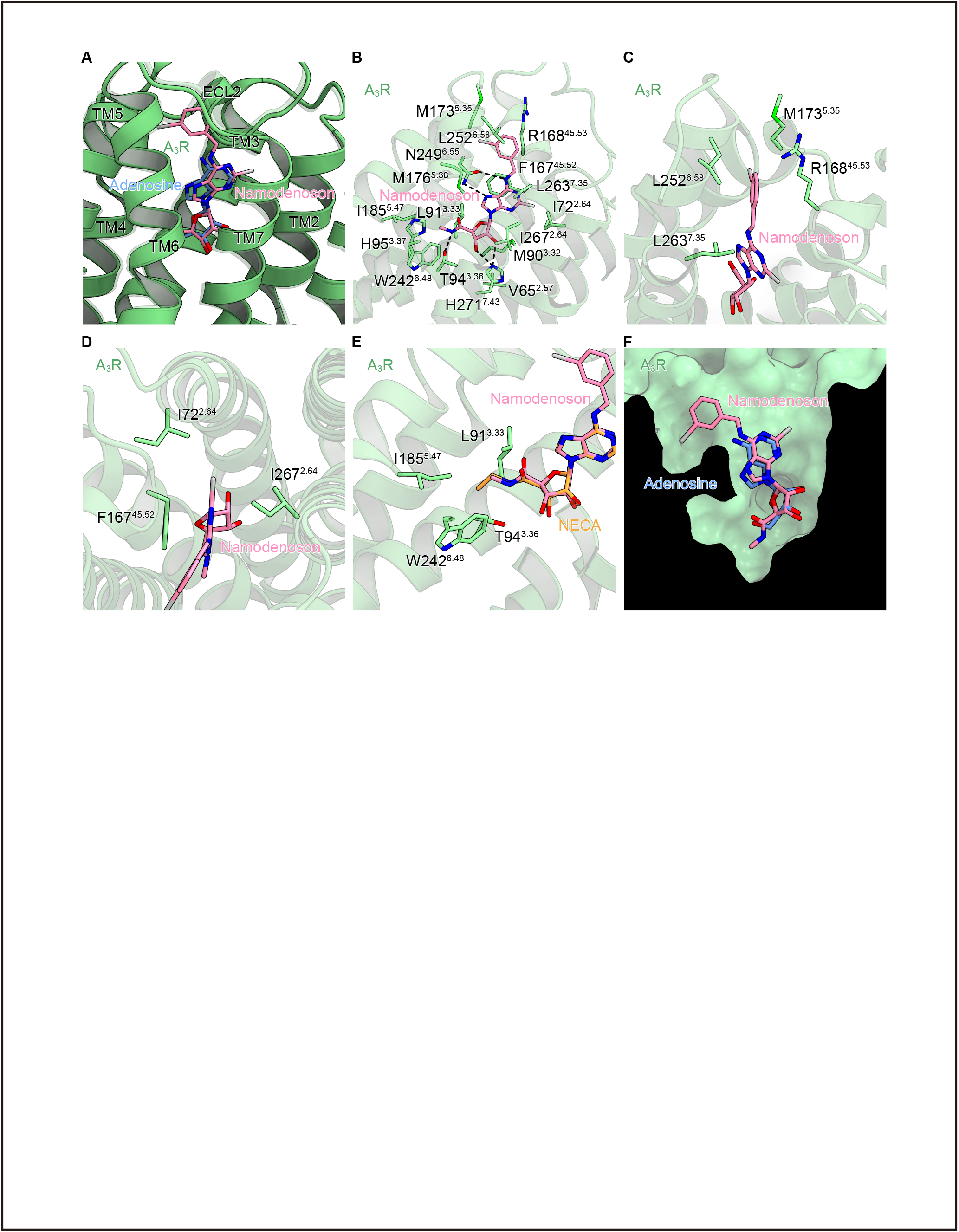
Binding mode of namodenoson. (A) Superimposition of adenosine- and namodenoson-bound A_3_R. The adenosine-bound A_3_R is represented by a translucent ribbon model. (B) Binding mode of namodenoson. (C) Close-up view of the hydrophobic pocket of the namodenoson-bound structure. The 3-iodobenzyl group of namodenoson wedges into the hydrophobic pocket. (D) Close-up view of the interactions around the chloro group of namodenoson. (E) Close-up view of the interactions around the methylamino group of namodenoson. (F) Cross section of the orthosteric pocket of A_3_R. Namodenoson fills the orthosteric pocket comprehensively.

In addition to the modification at the *N*^6^ position, that at the 2 position reportedly enhances the potency for A_3_R(Kim et al., 1994). The chloro group at the 2 position enters a cavity formed by TM1, TM2, and TM7, forming hydrophobic interactions with I72^2.64^ and I267^7.39^ (Figure 6D). These additional hydrophobic interactions should further stabilize the ligand-receptor interaction. Like the 5’ tail of NECA, the methylamino group at the 5’ position of namodenoson forms additional interactions, such as a hydrogen bond with T94^3.36^ (Figure 6E). However, compared to NECA, the 5’ tail of namodenoson is shorter, resulting in a weaker hydrophobic interaction with W242^6.48^, which is closely related to the activation of a class A GPCR. This length might be optimal to efficiently activate A_3_R, while preventing the nonselective activation of the other adenosine receptors. This inference is consistent with the report that compounds with a methylamino group at the 5’ position exhibited higher selectivity for A_3_R than those with an ethylamino group(Gallo-Rodriguez et al., 1994).

Overall, namodenoson exhibited unparalleled potency for A_3_R among the agonists (Figure S1C; Table S2). Compared to the other agonists, namodenoson fills the orthosteric pocket of A_3_R much more comprehensively (Figure 6F). Thus, each of the modifications at the *N*^6^, 2 and 5’ positions contributes to further the extensive and strong interactions, endowing namodenoson with its exceptional potency (EC_50_ = 2.41 nM for human A_3_R) and selectivity for A_3_R.

## Discussion

In the present study, we discovered novel modified adenosines that selectively activate A_3_R. All of these modified adenosines, including m^6^A, have hydrophobic modifications at the *N*^6^ position. We elucidated the structures of A_3_R complexes bound to two nonselective and three selective agonists including an A_3_R-selective agonist in clinical trials: namodenoson. While the binding modes of the adenosine moieties of these ligands are similar, remarkable differences were observed in the interactions between their modifications and the receptor. According to our model, we examined the ligand binding profiles of a comprehensive set of hydrophobic pocket mutants. Based on the results, we created A_3_R mutants that selectively inhibit the m^6^A pathway. Our structures and functional mutants will greatly aid drug discovery and molecular biological research targeting A_3_R and modified adenosines.

It should be noted that while unmodified adenosine can be generated by multiple pathways, including anabolic purine synthesis or the catabolic breakdown of ATP or RNA, m^6^A and i^6^A are exclusively generated through breakdown of RNA. Our previous study has shown that extracellular m^6^A is mostly derived from lysosomal degradation of ribosome RNA, and extracellular release of m^6^A is augmented by cytotoxic stimulation, leading to A_3_R-dependent inflammation and allergy in vivo. Our cryo-EM analysis thus provides the molecular basis underlying m^6^A-mediated immune response. In addition, the m^6^A-selective A_3_R mutant generated in this study paves a way to functionally discriminate the impact of adenosine-dependent and m^6^A-dependent A_3_R signaling in cells and in vivo.

In addition to m^6^A, we discovered that i^6^A is a novel endogenous ligand of A_3_R. i^6^A is exclusively found in tRNA including cytosolic tRNA^Sec^ and tRNA^Ser^, mitochondrial tRNA^Ser(UCN)^, tRNA^Trp^, tRNA^Tyr^, tRNA^Phe^. It is most likely that the extracellular i^6^A is derived from the breakdown of these tRNAs, however, the catabolism pathway needs to be elucidated in the future study. In addition, the pathophysiological role of i^6^A-mediated A_3_R activation and the functional difference with m^6^A also need to be clarified. Curiously, tRNA-derived i^6^A and its derivatives can serve as growth hormones through interaction with three cytokinin receptors, which are known as Arabidopsis Histidine Kinase (AHK) 2/3/4. AHK2/3/4 are highly conserved in plants but not in animals. Furthermore, there is no structural similarity between A_3_R and AHKs, suggesting that i^6^A might acquire a distinct function during the evolution. Based on our structural analysis and mutagenesis study, we discovered some mutations in the hydrophobic pockets that can greatly suppress adenosine and m^6^A activity but in turn enhanced i^6^A-mediated A_3_R activation. Future study using these mutant A_3_R will elucidate the pathological function of i^6^A-mediated A_3_R action.

Previous SAR studies showed that the additional hydrophobic modifications at the *N*^6^ position significantly enhanced the A_3_R selectivity(Ciancetta and Jacobson, 2017; “(N)-Methanocarba 2,N6-Disubstituted Adenine Nucleosides as Highly Potent and Selective A3 Adenosine Receptor Agonists | Journal of Medicinal Chemistry,” n.d.; Ohno et al., 2004). Along with namodenoson, agents such as piclidenoson, MRS5980, MRS5698, and FM101 selectively activate A_3_R(Fishman, 2022; Jacobson et al., 2019; Jeong et al., 2008). All of these agonists have a hydrophobic modification at the *N*^6^ position of the adenosine moiety, suggesting that they achieve their A_3_R selectivity through interactions similar to those observed between namodenoson and A_3_R. Some A_3_R agonists have a larger hydrophobic modification at the 2 position. Compared to namodenoson, these ligands may have more extensive interactions with the cavity formed by TM1, TM2, and TM7.

To comprehensively understand the selective activation of adenosine receptors, we also compared the structures of adenosine receptors bound to their selective drugs(Chen et al., 2022; Doré et al., 2011; Glukhova et al., 2017; Xu et al., 2011) (Figures S5A-S5F). While occupying the orthosteric pocket, each ligand exhibits a unique extension on the extracellular side. The modification at the 2 position of namodenoson interacts with residues in TM2 and TM7 of A_3_R, whereas its modification at the *N*^6^ position, like m^6^A, interacts with ECL2 and the extracellular sides of TM5, TM6 and TM7 (Figure S5A). In contrast to namodenoson, the modification of i^6^A interacts only with the N-terminal side of ECL2 of A_3_R and the extracellular side of TM7 (Figure S5B). dU172, an A_1_R-selective agonist, not only interacts with the extracellular sides of TM5, TM6, and TM7, but also penetrates the cavity formed by TM1, TM2, and TM7 of A_1_R (Figure S5C). This insertion into the cavity formed by TM1, TM2, and TM7 also occurred with ZM241385-A_2A_R and BAY60-6583-A_2B_R (Figures S5D and S5E), indicating that this cavity is a major region contributing to the selectivity of adenosine receptors. Furthermore, the A_2A_R-selective agonist UK432097 exhibits a substantial extension of its modification site on the extracellular side, engaging in numerous interactions with TM2, ECL2, TM6, TM7, and ECL3 (Figure S5F). As observed in the sequence alignment of the adenosine receptors, while the “insides” of a ligand pocket are remarkably similar, unique interaction sites existing “outside” of the pocket have relatively diverse sequences (Figure S3). Thus, the selective activation of an adenosine receptor may be characterized by two factors: nonselective binding in the adenosine-binding region and selective binding on the extracellular side. In this study, we present the newly identified, unique hydrophobic pocket of A_3_R, which is not involved in ligand-receptor interactions in the other adenosine receptors. This structural insight will greatly facilitate the rational design of selective drugs targeting A_3_R as well as the other adenosine receptors.

## Methods

### Screening of A_3_R-selective nucleosides by TGFα-shedding assay

To measure the activation of the adenosine receptor subtypes (A_1_R, A_2A_R, A_2B_R, A_3_R), a transforming growth factor-α (TGFα) shedding assay was performed(Inoue et al., 2012). Briefly, HEK293A cells were seeded in 6-cm culture dishes. Cells were transfected using a polyethylenimine (PEI) agent (10 μl of 1 mg/ml per dish hereafter; Polysciences) with pCAG plasmids encoding human A_1_R, A_2A_R, A_2B_R, A_3_R, or mock (400 ng per dish hereafter), together with the plasmids encoding alkaline phosphatase (AP)-tagged TGFα (AP-TGFα; 1 µg) and chimeric Gα subunit proteins (Gαq/o for A_1_R, Gαq/s for A_2A_R and A_2B_R, and Gα_q/i3_ for A_3_R; 200 ng). These plasmids were gifts from A. Inoue, Tohoku University. After 24 h culture, the transfected cells were harvested using trypsin-EDTA, neutralized with FBS-containing DMEM and collected by centrifugation. Cells were resuspended in Hank’s Balanced Salt Solution (HBSS, GIBCO) containing 5 mM HEPES (pH 7.4) and seeded in a 96-well plate. After 30 min culture, each nucleoside was adjusted with 0.01% bovine serum albumin (BSA)-containing HEPES-HBSS to a concentration comparable to the EC_50_ of unmodified adenosine for each receptor(Ogawa et al., 2021) and then added to the cells (final concentrations of 100 nM for A_1_R, 1 µM for A_2A_R, 5 µM for A_2B_R and 100 nM for A_3_R). After a 1 h incubation, the conditioned media were transferred to an empty 96-well plate. The reaction solution (a mixture of 10 mM p-nitrophenylphosphate (p-NPP), 120mM Tris-HCl (pH 9.5), 40 mM NaCl, and 10 mM MgCl2) was added to plates containing cells and conditioned media. The absorbance at a wavelength of 405 nm was measured using a microplate reader (Molecular Devices) before and after a 1-h incubation of the plates at room temperature. AP-TGFα release was calculated by subtracting the spontaneous AP-TGFα release signal from the nucleoside-induced AP-TGFα release signal. Details of each tested nucleoside are provided in Table S4.

### Expression and purification of A_3_R-G_i_ complex

The gene encoding full-length sheep A_3_R (Uniprot ID: W5QED6) was constructed with the native signal peptide replaced with the HA-signal peptide and the DYKDDDDK Flag epitope tag. LgBiT was fused to the C-terminus of A_3_R, followed by a 3C protease site and an eGFP-His8 tag. GGSGGGGSGGSSSGG linkers were inserted on both the N-terminal and C-terminal sides of LgBiT. Human G_i1_ was fused to the C-terminus of bovine G_γ2_, following the GSAGSAGSA linker. Rat G_β1_ was cloned with a C-terminal HiBiT connected with the 15 amino sequence GGSGGGGSGGSSSGG. The resulting constructs were subcloned into the pEG BacMam vector.

The viruses expressing A_3_R, G_γ2_-G_i1_, and G_β1_-HiBiT were prepared using the BacMam system. A liter of HEK293S GnTI (N-acetylglucosaminyl-transferase I–negative) cells at a concentration of 3 × 10^6^ cells/mL was co-infected with the prepared viruses expressed at the ratio of 1:1:1.

The collected cells were resuspended and Dounce-homogenized in 20 mM Tris-HCl, pH 8.0, 100 mM NaCl, 10 % glycerol, and 10 μM ligand. After homogenization, 25 mU/mL of apyrase and 10 μM of agonist were added and the lysate was rotated at room temperature for 1 hr. Afterwards, the membrane fraction was isolated through ultracentrifugation at 180,000*g* for 1 h, and solubilized in 20 mM Tris-HCl, pH 8.0, 150 mM NaCl, 1 % DDM, 0.2 % CHS, 10 % glycerol, and 10 μM ligand, at 4 °C for 1 hr. The insoluble fraction was removed by ultracentrifugation at 180,000*g* for 30 min and the supernatant was then incubated with Anti-DYKDDDDK M1 resin (Genscript) for 1 h. The resin was washed with 20 column volumes of buffer, containing 20 mM Tris-HCl, pH 8.0, 500 mM NaCl, 10 % glycerol, 0.1 % Lauryl Maltose Neopentyl Glycol (LMNG; Anatrace), 0.01 % CHS, and 1-10 μM ligand. The complex was eluted with buffer, containing 20 mM Tris-HCl, pH 8.0, 150 mM NaCl, 10 % glycerol, 0.1 % LMNG, 0.01 % CHS, 1-10 μM ligand, and 0.15 mg/ml Flag peptide. The eluate was incubated with 0.5 mg of HRV-3C protease (prepared in-house) and 0.5 mg of scFv16 (prepared as described previously(Okamoto et al., 2021)) at 4 °C overnight. The complex was concentrated and separated by size exclusion chromatography on a Superose 6 Increase 10/300 column, in buffer containing 20 mM Tris-HCl, pH 8.0, 150 mM NaCl, 0.01 % LMNG, 0.001 % CHS, and 1-10 μM agonist. Peak fractions were concentrated for the cryo-EM grid preparation.

### Cryo-EM data acquisition

A 3 μL portion of the purified complex was deposited onto a Quantifoil holey carbon grid (R1.2/1.3, Au, 300 mesh), which had been glow-discharged just before use. This grid was then rapidly frozen in liquid ethane using a Vitrobot Mark IV (FEI). Cryo-EM data acquisition was performed with a Titan Krios G3i microscope (300 kV, Thermo Fisher Scientific), outfitted with a BioQuantum K3 imaging filter (Gatan) and a K3 direct electron detector (Gatan). Movies, each with a calibrated pixel size of 0.83 Å per pixel and a defocus range between −0.8 and −1.6 μm, were acquired utilizing Thermo Fisher Scientific’s EPU software for single-particle data collection. Movies were recorded over 2.3 s and divided into 48 frames. The numbers and total exposures of movies are provided in Table 1.

**Table. 1.** Cryo-EM.

| **Data collection** | NECA-A<sub>3</sub>R-G<sub>i</sub> | adenosine-A<sub>3</sub>R-G<sub>i</sub> | m<sup>6</sup>A-A<sub>3</sub>R-G<sub>i</sub> | i<sup>6</sup>A-A<sub>3</sub>R-G<sub>i</sub> | namodenoson-A<sub>3</sub>R-G<sub>i</sub> |
| --- | --- | --- | --- | --- | --- |
| Microscope | Titan Krios (Thermo Fisher Scientific) |  |  |  |  |
| Voltage (keV) | 300 |  |  |  |  |
| Electron exposure (e<sup>-</sup>/Å<sup>2</sup>) | 48 | 48.30 | 50.062 | 49.998 | 50 |
| Detector | Gatan K3 Summit camera (Gatan) |  |  |  |  |
| Magnification | ×105,000 |  |  |  |  |
| Defocus range (μm) | -0.8–1.6 |  |  |  |  |
| Pixel size (Å/pix) | 0.83 |  |  |  |  |
| Number of movies | 15,004 | 21,810 | 12,663 | 21,300 | 20,609 |
| Symmetry | C1 |  |  |  |  |
| Picked particles | 5,682,938 | 6,981,460 | 3,200,082 | 17,004,284 | 15,299,681 |
| Final particles | 210,192 | 160,357 | 144,012 | 141,536 | 65,161 |
| Map resolution (Å) | 2.86 | 3.27 | 3.40 | 3.66 | 3.62 |
| FSC threshold | 0.143 |  |  |  |  |
| **Model refinement** |  |  |  |  |  |
| Atoms | 17339 | 17377 | 17371 | 17366 | 17390 |
| **R.m.s. deviations from ideal** |  |  |  |  |  |
| Bond lengths (Å) | 0.003 | 0.003 | 0.003 | 0.003 | 0.004 |
| Bond angles (°) | 0.63 | 0.537 | 0.509 | 0.567 | 0.613 |
| Validation |  |  |  |  |  |
| Clashscore | 8.39 | 11.99 | 10.38 | 14.70 | 20.34 |
| Rotamers (%) | 0.00 | 0.00 | 0.00 | 0.00 | 0.10 |
| **Ramachandran plot** |  |  |  |  |  |
| Favored (%) | 94.03 | 95.21 | 95.94 | 95.12 | 93.82 |
| Allowed (%) | 5.97 | 4.70 | 3.96 | 4.79 | 6.08 |
| Outlier (%) | 0 | 0.09 | 0.09 | 0.09 | 0.09 |

The movies, captured in super-resolution mode, were processed by binning them 2×, followed by dose fractionation and correction for beam-induced motion using RELION 4.0 or cryoSPARC v4.0(Punjani et al., 2017; Zivanov et al., 2018). The contrast transfer function (CTF) parameters were estimated using patch CTF estimation in cryoSPARC.

Although the workflow for each complex differs in its details, the outlines are similar. Thus, the workflow for the NECA-bound complex is described here. The workflows for all complexes are summarized in Figures S2A-S2E.

A subset of particles was initially identified using the Blob picker from a selection of micrographs, and then subjected to multiple rounds of 2D classification in cryoSPARC. Selected particles were used for training the Topaz model(Bepler et al., 2019). For the full dataset, 5,682,938 particles were picked and extracted. Subsequent rounds of hetero-refinement were performed to discard poor-quality particle classes. The refined particles were further refined through 3D classification without alignment in RELION. This process yielded 210,192 high-quality particles in the optimal class, which were reconstructed using non-uniform refinement in cryoSPARC to achieve an overall resolution of 2.86 Å, based on the gold standard Fourier Shell Correlation (FSC = 0.143) criteria. Afterwards, the 3D model was refined with masks on the receptor and the G protein trimer. The locally refined maps were integrated using phenix.combine_focused_maps for model construction(Liebschner et al., 2019).

### Model building and refinement

The AlphaFold2-predicted sheep A_3_R model and the cryo-EM structure of the MT_1_-G_i1_ complex (PDB ID: 7DB6) were used as starting templates for modeling the A_3_R and G_i1_ components of the NECA-bound structure, respectively(Jumper et al., 2021; Okamoto et al., 2021). Initially, these models were positioned into the density map using jiggle fit in COOT, and subsequently, the atomic models were fine-tuned using COOT and refined with phenix.real_space_refine (v1.19), incorporating secondary structure restraints from phenix.secondary_structure_restraints. Restraints for ligands were generated with Grade2 (Smart, O.S., Sharff A., Holstein, J., Womack, T.O., Flensburg, C., Keller, P., Paciorek, W., Vonrhein, C. and Bricogne G. (2021) Grade2 version 1.5.0. Cambridge, United Kingdom: Global Phasing Ltd.). The NECA-bound model was used as the initial model for the other complexes, while the following procedures were the same as those used for the NECA-bound structure.

### Mutagenesis study to investigate the specificity of the modified adenosines

A total of 34 A_3_R mutants for V169^45.53^, I253^5.35^, M174^6.58^, and L264^7.35^ were generated by introducing single point mutations into pCAG plasmids encoding human A_3_R, using a KOD-Plus-Mutagenesis kit (TOYOBO), and confirmed by Sanger sequencing (Azenta). These plasmids were transfected into cells using a PEI agent with AP-TGFα and Gαq/i3, in the same volume as mentioned above, and the TGFα-shedding assay was performed as described above. Unmodified adenosine, m^6^A, m^6,6^A, and i^6^A were 10-fold serially diluted with 0.01% BSA-containing HEPES-HBSS to give final concentrations in the range of 100 µM to 1 nM. AP-TGFα release percentages were fitted to a four-parameter sigmoidal concentration–response curve, using the Prism 9 software (GraphPad Prism), and the EC_50_ and E_max_ values were obtained. Receptor activation is scored using the relative intrinsic activity (RAi), which is defined as the relative E_max_/ EC_50_ value(Ehlert et al., 1999; Inoue et al., 2012).

## Supporting information

Supplementary items

## Data Availability

Atomic coordinates for the NECA-A3R-Gi, adenosine-A3R-Gi, m6A-A3R-Gi, namodenoson-A3R-Gi, i6A-A3R-Gi complex have been deposited in the Protein Data Bank, under accession code 8YH0, 8YH2, 8YH3, 8YH6, and 8YH5, respectively. The associated electron microscopy data have been deposited in the Electron Microscopy Database and EMD-39278, EMD-39279, EMD-39280, EMD-39282, and EMD-39281, respectively. All other data are available from the corresponding author upon reasonable request.

## Acknowledgments

We thank K. Ogomori and C. Harada for technical assistance. This work was funded by the JSPS KAKENHI (Grant Nos. 21H05037 to O.N., 22K19371 and 22H02751 to W.S., JP21H02659 and JP21H05265 to F.-Y.W.), the ONO Medical Research Foundation, the Kao Foundation for Arts and Sciences, The Uehara Memorial Foundation (all to W.S.), the Takeda Science Foundation (to W.S. and F.-Y.W.), the FOREST program from JST (Grant No. JPMJFR220K to A.O.), the ERATO program from JST (Grant No. JPMJER2002 to F.Y.W.), and the AMED Platform Project for Supporting Drug Discovery and Life Science Research (BINDS) (Grant No. JP19am01011115, support No. 1109 to O.N.).

## Author contributions

H.S.O. performed the complex purification, the cryo-EM data collection, the single particle analysis, model building and refinement, and designed the mutants with assistance from F.K.S., H.A., H.H.O., A.I., C.N., and W.S. A.O. performed the screening and the cell-based assays, with assistance from T.K. and F.-Y.W. H.S.O., A.O., F.-Y.W, W.S., O.N. wrote the manuscript with inputs from all authors. F.-Y.W., W.S., and O.N. supervised the research.

## References

Akasaka, H., Tanaka, T., Sano, F.K., Matsuzaki, Y., Shihoya, W., Nureki, O., 2022. Structure of the active Gi-coupled human lysophosphatidic acid receptor 1 complexed with a potent agonist. Nat. Commun. 13, 5417. 10.1038/s41467-022-33121-2

Ballesteros, J.A., Weinstein, H., 1995. [19] Integrated methods for the construction of three-dimensional models and computational probing of structure-function relations in G protein-coupled receptors, in: Sealfon, S.C. (Ed.), Methods in Neurosciences, Receptor Molecular Biology. Academic Press, pp. 366–428. 10.1016/S1043-9471(05)80049-7

Bepler, T., Morin, A., Rapp, M., Brasch, J., Shapiro, L., Noble, A.J., Berger, B., 2019. Positive-unlabeled convolutional neural networks for particle picking in cryo-electron micrographs. Nat. Methods 16, 1153–1160. 10.1038/s41592-019-0575-8

Cai, H., Xu, Y., Guo, S., He, X., Sun, J., Li, X., Li, C., Yin, W., Cheng, X., Jiang, H., Xu, H.E., Xie, X., Jiang, Y., 2022. Structures of adenosine receptor A2BR bound to endogenous and synthetic agonists. Cell Discov. 8, 1–4. 10.1038/s41421-022-00503-1

Carpenter, B., Nehmé, R., Warne, T., Leslie, A.G.W., Tate, C.G., 2016. Structure of the adenosine A2A receptor bound to an engineered G protein. Nature 536, 104–107. 10.1038/nature18966

Chen, Y., Zhang, J., Weng, Y., Xu, Y., Lu, W., Liu, W., Liu, M., Hua, T., Song, G., 2022. Cryo-EM structure of the human adenosine A2B receptor–Gs signaling complex. Sci. Adv. 8, eadd3709. 10.1126/sciadv.add3709

Cheng, R.K.Y., Segala, E., Robertson, N., Deflorian, F., Doré, A.S., Errey, J.C., Fiez-Vandal, C., Marshall, F.H., Cooke, R.M., 2017. Structures of Human A1 and A2A Adenosine Receptors with Xanthines Reveal Determinants of Selectivity. Structure 25, 1275–1285.e4. 10.1016/j.str.2017.06.012

Ciancetta, A., Jacobson, K.A., 2017. Structural Probing and Molecular Modeling of the A3 Adenosine Receptor: A Focus on Agonist Binding. Molecules 22, 449. 10.3390/molecules22030449

Cieślak, M., Komoszyński, M., Wojtczak, A., 2008. Adenosine A2A receptors in Parkinson’s disease treatment. Purinergic Signal. 4, 305–312. 10.1007/s11302-008-9100-8

Doré, A.S., Robertson, N., Errey, J.C., Ng, I., Hollenstein, K., Tehan, B., Hurrell, E., Bennett, K., Congreve, M., Magnani, F., Tate, C.G., Weir, M., Marshall, F.H., 2011. Structure of the Adenosine A2A Receptor in Complex with ZM241385 and the Xanthines XAC and Caffeine. Structure 19, 1283–1293. 10.1016/j.str.2011.06.014

Draper-Joyce, C.J., Bhola, R., Wang, J., Bhattarai, A., Nguyen, A.T.N., Cowie-Kent, I., O’Sullivan, K., Chia, L.Y., Venugopal, H., Valant, C., Thal, D.M., Wootten, D., Panel, N., Carlsson, J., Christie, M.J., White, P.J., Scammells, P., May, L.T., Sexton, P.M., Danev, R., Miao, Y., Glukhova, A., Imlach, W.L., Christopoulos, A., 2021. Positive allosteric mechanisms of adenosine A1 receptor-mediated analgesia. Nature 597, 571–576. 10.1038/s41586-021-03897-2

Draper-Joyce, C.J., Khoshouei, M., Thal, D.M., Liang, Y.-L., Nguyen, A.T.N., Furness, S.G.B., Venugopal, H., Baltos, J.-A., Plitzko, J.M., Danev, R., Baumeister, W., May, L.T., Wootten, D., Sexton, P.M., Glukhova, A., Christopoulos, A., 2018. Structure of the adenosine-bound human adenosine A1 receptor–Gi complex. Nature 558, 559–563. 10.1038/s41586-018-0236-6

Duan, J., Shen, D.-D., Zhou, X.E., Bi, P., Liu, Q.-F., Tan, Y.-X., Zhuang, Y.-W., Zhang, H.-B., Xu, P.-Y., Huang, S.-J., Ma, S.-S., He, X.-H., Melcher, K., Zhang, Y., Xu, H.E., Jiang, Y., 2020. Cryo-EM structure of an activated VIP1 receptor-G protein complex revealed by a NanoBiT tethering strategy. Nat. Commun. 11, 4121. 10.1038/s41467-020-17933-8

Effendi, W.I., Nagano, T., Kobayashi, K., Nishimura, Y., 2020. Focusing on Adenosine Receptors as a Potential Targeted Therapy in Human Diseases. Cells 9, 785. 10.3390/cells9030785

Ehlert, F.J., Griffin, M.T., Sawyer, G.W., Bailon, R., 1999. A simple method for estimation of agonist activity at receptor subtypes: comparison of native and cloned M3 muscarinic receptors in guinea pig ileum and transfected cells. J. Pharmacol. Exp. Ther. 289, 981–992.

Fishman, P., 2022. Drugs Targeting the A3 Adenosine Receptor: Human Clinical Study Data. Molecules 27, 3680. 10.3390/molecules27123680

Flock, T., Ravarani, C.N.J., Sun, D., Venkatakrishnan, A.J., Kayikci, M., Tate, C.G., Veprintsev, D.B., Babu, M.M., 2015. Universal allosteric mechanism for Gα activation by GPCRs. Nature 524, 173–179. 10.1038/nature14663

Fredholm, B.B., IJzerman, A.P., Jacobson, K.A., Klotz, K.N., Linden, J., 2001. International Union of Pharmacology. XXV. Nomenclature and classification of adenosine receptors. Pharmacol. Rev. 53, 527–552.

Gallo-Rodriguez, C., Ji, X.D., Melman, N., Siegman, B.D., Sanders, L.H., Orlina, J., Fischer, B., Pu, Q., Olah, M.E., van Galen, P.J., 1994. Structure-activity relationships of N6-benzyladenosine-5’-uronamides as A3-selective adenosine agonists. J. Med. Chem. 37, 636–646. 10.1021/jm00031a014

Gallos, G., Ruyle, T.D., Emala, C.W., Lee, H.T., 2005. A1 adenosine receptor knockout mice exhibit increased mortality, renal dysfunction, and hepatic injury in murine septic peritonitis. Am. J. Physiol. Renal Physiol. 289, F369–376. 10.1152/ajprenal.00470.2004

Gao, Z.-G., Auchampach, J.A., Jacobson, K.A., 2023. Species dependence of A3 adenosine receptor pharmacology and function. Purinergic Signal. 19, 523–550. 10.1007/s11302-022-09910-1

García-Nafría, J., Lee, Y., Bai, X., Carpenter, B., Tate, C.G., 2018. Cryo-EM structure of the adenosine A2A receptor coupled to an engineered heterotrimeric G protein. eLife 7, e35946. 10.7554/eLife.35946

Gessi, S., Merighi, S., Varani, K., Leung, E., Mac Lennan, S., Borea, P.A., 2008. The A3 adenosine receptor: An enigmatic player in cell biology. Pharmacol. Ther. 117, 123–140. 10.1016/j.pharmthera.2007.09.002

Glukhova, A., Thal, D.M., Nguyen, A.T., Vecchio, E.A., Jörg, M., Scammells, P.J., May, L.T., Sexton, P.M., Christopoulos, A., 2017. Structure of the Adenosine A1 Receptor Reveals the Basis for Subtype Selectivity. Cell 168, 867–877.e13. 10.1016/j.cell.2017.01.042

Guo, D., Pan, A.C., Dror, R.O., Mocking, T., Liu, R., Heitman, L.H., Shaw, D.E., IJzerman, A.P., 2016. Molecular Basis of Ligand Dissociation from the Adenosine A2A Receptor. Mol. Pharmacol. 89, 485–491. 10.1124/mol.115.102657

Hattori, M., Hibbs, R.E., Gouaux, E., 2012. A fluorescence-detection size-exclusion chromatography-based thermostability assay for membrane protein precrystallization screening. Struct. Lond. Engl. 1993 20, 1293–1299. 10.1016/j.str.2012.06.009

Hua, T., Li, X., Wu, L., Iliopoulos-Tsoutsouvas, C., Wang, Y., Wu, M., Shen, L., Brust, C.A., Nikas, S.P., Song, F., Song, X., Yuan, S., Sun, Q., Wu, Y., Jiang, S., Grim, T.W., Benchama, O., Stahl, E.L., Zvonok, N., Zhao, S., Bohn, L.M., Makriyannis, A., Liu, Z.-J., 2020. Activation and Signaling Mechanism Revealed by Cannabinoid Receptor-Gi Complex Structures. Cell 180, 655–665.e18. 10.1016/j.cell.2020.01.008

Inoue, A., Ishiguro, J., Kitamura, H., Arima, N., Okutani, M., Shuto, A., Higashiyama, S., Ohwada, T., Arai, H., Makide, K., Aoki, J., 2012. TGFα shedding assay: an accurate and versatile method for detecting GPCR activation. Nat. Methods 9, 1021–1029. 10.1038/nmeth.2172

Iwama, A., Akasaka, H., Sano, F.K., Oshima, H.S., Shihoya, W., Nureki, O., 2023. Structure and dynamics of the RF-amide QRFP receptor GPR103. 10.1101/2023.12.06.570340

Izume, T., Kawahara, R., Uwamizu, A., Chen, L., Yaginuma, S., Omi, J., Kawana, H., Hou, F., Sano, F.K., Tanaka, T., Kobayashi, K., Okamoto, H.H., Kise, Y., Ohwada, T., Aoki, J., Shihoya, W., Nureki, O., 2024. Structural basis for lysophosphatidylserine recognition by GPR34. Nat. Commun. 15, 902. 10.1038/s41467-024-45046-z

Jaakola, V.-P., Griffith, M.T., Hanson, M.A., Cherezov, V., Chien, E.Y.T., Lane, J.R., IJzerman, A.P., Stevens, R.C., 2008. The 2.6 Angstrom Crystal Structure of a Human A2A Adenosine Receptor Bound to an Antagonist. Science 322, 1211–1217. 10.1126/science.1164772

Jacobson, K.A., Tosh, D.K., Jain, S., Gao, Z.-G., 2019. Historical and Current Adenosine Receptor Agonists in Preclinical and Clinical Development. Front. Cell. Neurosci. 13, 124. 10.3389/fncel.2019.00124

Jeong, L.S., Pal, S., Choe, S.A., Choi, W.J., Jacobson, K.A., Gao, Z.-G., Klutz, A.M., Hou, X., Kim, H.O., Lee, H.W., Lee, S.K., Tosh, D.K., Moon, H.R., 2008. Structure-activity relationships of truncated D- and l-4’-thioadenosine derivatives as species-independent A3 adenosine receptor antagonists. J. Med. Chem. 51, 6609–6613. 10.1021/jm8008647

Jumper, J., Evans, R., Pritzel, A., Green, T., Figurnov, M., Ronneberger, O., Tunyasuvunakool, K., Bates, R., Žídek, A., Potapenko, A., Bridgland, A., Meyer, C., Kohl, S.A.A., Ballard, A.J., Cowie, A., Romera-Paredes, B., Nikolov, S., Jain, R., Adler, J., Back, T., Petersen, S., Reiman, D., Clancy, E., Zielinski, M., Steinegger, M., Pacholska, M., Berghammer, T., Bodenstein, S., Silver, D., Vinyals, O., Senior, A.W., Kavukcuoglu, K., Kohli, P., Hassabis, D., 2021. Highly accurate protein structure prediction with AlphaFold. Nature 596, 583–589. 10.1038/s41586-021-03819-2

Kato, H.E., Zhang, Y., Hu, H., Suomivuori, C.-M., Kadji, F.M.N., Aoki, J., Krishna Kumar, K., Fonseca, R., Hilger, D., Huang, W., Latorraca, N.R., Inoue, A., Dror, R.O., Kobilka, B.K., Skiniotis, G., 2019. Conformational transitions of a neurotensin receptor 1–Gi1 complex. Nature 572, 80–85. 10.1038/s41586-019-1337-6

Kim, H.O., Ji, X.D., Siddiqi, S.M., Olah, M.E., Stiles, G.L., Jacobson, K.A., 1994. 2-Substitution of N6-benzyladenosine-5’-uronamides enhances selectivity for A3 adenosine receptors. J. Med. Chem. 37, 3614–3621. 10.1021/jm00047a018

Kim, K., Che, T., Panova, O., DiBerto, J.F., Lyu, J., Krumm, B.E., Wacker, D., Robertson, M.J., Seven, A.B., Nichols, D.E., Shoichet, B.K., Skiniotis, G., Roth, B.L., 2020. Structure of a Hallucinogen-Activated Gq-Coupled 5-HT2A Serotonin Receptor. Cell 182, 1574–1588.e19. 10.1016/j.cell.2020.08.024

Lebon, G., Warne, T., Edwards, P.C., Bennett, K., Langmead, C.J., Leslie, A.G.W., Tate, C.G., 2011. Agonist-bound adenosine A2A receptor structures reveal common features of GPCR activation. Nature 474, 521–525. 10.1038/nature10136

Liebschner, D., Afonine, P.V., Baker, M.L., Bunkóczi, G., Chen, V.B., Croll, T.I., Hintze, B., Hung, L.-W., Jain, S., McCoy, A.J., Moriarty, N.W., Oeffner, R.D., Poon, B.K., Prisant, M.G., Read, R.J., Richardson, J.S., Richardson, D.C., Sammito, M.D., Sobolev, O.V., Stockwell, D.H., Terwilliger, T.C., Urzhumtsev, A.G., Videau, L.L., Williams, C.J., Adams, P.D., 2019. Macromolecular structure determination using X-rays, neutrons and electrons: recent developments in Phenix. Acta Crystallogr. Sect. Struct. Biol. 75, 861–877. 10.1107/S2059798319011471

Liu, H., Zeng, T., He, C., Rawal, V.H., Zhou, H., Dickinson, B.C., 2022. Development of mild chemical catalysis conditions for m1A-to-m6A rearrangement on RNA. ACS Chem. Biol. 17, 1334. 10.1021/acschembio.2c00178

Madi, L., Ochaion, A., Rath-Wolfson, L., Bar-Yehuda, S., Erlanger, A., Ohana, G., Harish, A., Merimski, O., Barer, F., Fishman, P., 2004. The A3 adenosine receptor is highly expressed in tumor versus normal cells: potential target for tumor growth inhibition. Clin. Cancer Res. Off. J. Am. Assoc. Cancer Res. 10, 4472–4479. 10.1158/1078-0432.CCR-03-0651

Nagiri, C., Kobayashi, Kazuhiro, Tomita, A., Kato, M., Kobayashi, Kan, Yamashita, K., Nishizawa, T., Inoue, A., Shihoya, W., Nureki, O., 2021. Cryo-EM structure of the β3-adrenergic receptor reveals the molecular basis of subtype selectivity. Mol. Cell 81, 3205–3215.e5. 10.1016/j.molcel.2021.06.024

(N)-Methanocarba 2,N6-Disubstituted Adenine Nucleosides as Highly Potent and Selective A3 Adenosine Receptor Agonists | Journal of Medicinal Chemistry [WWW Document], n.d. URL https://pubs.acs.org/doi/full/10.1021/jm049580r (accessed 2.15.24).

Nureki, I., Kobayashi, K., Tanaka, T., Demura, K., Inoue, A., Shihoya, W., Nureki, O., 2022. Cryo-EM structures of the β3 adrenergic receptor bound to solabegron and isoproterenol. Biochem. Biophys. Res. Commun. 611, 158–164. 10.1016/j.bbrc.2022.04.065

Ogawa, A., Nagiri, C., Shihoya, W., Inoue, A., Kawakami, K., Hiratsuka, S., Aoki, J., Ito, Y., Suzuki, Takeo, Suzuki, Tsutomu, Inoue, T., Nureki, O., Tanihara, H., Tomizawa, K., Wei, F.-Y., 2021. N6-methyladenosine (m6A) is an endogenous A3 adenosine receptor ligand. Mol. Cell 81, 659–674.e7. 10.1016/j.molcel.2020.12.038

Ohno, M., Gao, Z.-G., Van Rompaey, P., Tchilibon, S., Kim, S.-K., Harris, B.A., Gross, A.S., Duong, H.T., Van Calenbergh, S., Jacobson, K.A., 2004. Modulation of adenosine receptor affinity and intrinsic efficacy in adenine nucleosides substituted at the 2-position. Bioorg. Med. Chem. 12, 2995–3007. 10.1016/j.bmc.2004.03.031

Okamoto, H.H., Miyauchi, H., Inoue, A., Raimondi, F., Tsujimoto, H., Kusakizako, T., Shihoya, W., Yamashita, K., Suno, R., Nomura, N., Kobayashi, T., Iwata, S., Nishizawa, T., Nureki, O., 2021. Cryo-EM structure of the human MT1–Gi signaling complex. Nat. Struct. Mol. Biol. 28, 694–701. 10.1038/s41594-021-00634-1

Oshima, H.S., Sano, F.K., Akasaka, H., Iwama, A., Shihoya, W., Nureki, O., 2024. Optimizing cryo-EM structural analysis of Gi-coupling receptors via engineered Gt and Nb35 application. Biochem. Biophys. Res. Commun. 693, 149361. 10.1016/j.bbrc.2023.149361

Pándy-Szekeres, G., Munk, C., Tsonkov, T.M., Mordalski, S., Harpsøe, K., Hauser, A.S., Bojarski, A.J., Gloriam, D.E., 2018. GPCRdb in 2018: adding GPCR structure models and ligands. Nucleic Acids Res. 46, D440–D446. 10.1093/nar/gkx1109

Punjani, A., Rubinstein, J.L., Fleet, D.J., Brubaker, M.A., 2017. cryoSPARC: algorithms for rapid unsupervised cryo-EM structure determination. Nat. Methods 14, 290–296. 10.1038/nmeth.4169

Rasmussen, S.G.F., DeVree, B.T., Zou, Y., Kruse, A.C., Chung, K.Y., Kobilka, T.S., Thian, F.S., Chae, P.S., Pardon, E., Calinski, D., Mathiesen, J.M., Shah, S.T.A., Lyons, J.A., Caffrey, M., Gellman, S.H., Steyaert, J., Skiniotis, G., Weis, W.I., Sunahara, R.K., Kobilka, B.K., 2011. Crystal structure of the β2 adrenergic receptor–Gs protein complex. Nature 477, 549–555. 10.1038/nature10361

Robert, X., Gouet, P., 2014. Deciphering key features in protein structures with the new ENDscript server. Nucleic Acids Res. 42, W320–W324. 10.1093/nar/gku316

Sano, F.K., Akasaka, H., Shihoya, W., Nureki, O., 2023. Cryo-EM structure of the endothelin-1-ETB-Gi complex. eLife 12, e85821. 10.7554/eLife.85821

The UniProt Consortium, 2023. UniProt: the Universal Protein Knowledgebase in 2023. Nucleic Acids Res. 51, D523–D531. 10.1093/nar/gkac1052

Xu, F., Wu, H., Katritch, V., Han, G.W., Jacobson, K.A., Gao, Z.-G., Cherezov, V., Stevens, R.C., 2011. Structure of an Agonist-Bound Human A2A Adenosine Receptor. Science 332, 322–327. 10.1126/science.1202793

Yuan, Y., Jia, G., Wu, C., Wang, W., Cheng, L., Li, Q., Li, Z., Luo, K., Yang, S., Yan, W., Su, Z., Shao, Z., 2021. Structures of signaling complexes of lipid receptors S1PR1 and S1PR5 reveal mechanisms of activation and drug recognition. Cell Res. 31, 1263– 1274. 10.1038/s41422-021-00566-x

Zhang, K., Wu, H., Hoppe, N., Manglik, A., Cheng, Y., 2022. Fusion protein strategies for cryo-EM study of G protein-coupled receptors. Nat. Commun. 13, 4366. 10.1038/s41467-022-32125-2

Zhou, Q., Yang, D., Wu, M., Guo, Y., Guo, W., Zhong, L., Cai, X., Dai, A., Jang, W., Shakhnovich, E.I., Liu, Z.-J., Stevens, R.C., Lambert, N.A., Babu, M.M., Wang, M.-W., Zhao, S., 2019. Common activation mechanism of class A GPCRs. eLife 8, e50279. 10.7554/eLife.50279

Zhou, Q.Y., Li, C., Olah, M.E., Johnson, R.A., Stiles, G.L., Civelli, O., 1992. Molecular cloning and characterization of an adenosine receptor: the A3 adenosine receptor. Proc. Natl. Acad. Sci. 89, 7432–7436. 10.1073/pnas.89.16.7432

Zhuang, Y., Wang, Y., He, B., He, X., Zhou, X.E., Guo, S., Rao, Q., Yang, J., Liu, J., Zhou, Q., Wang, X., Liu, M., Liu, W., Jiang, X., Yang, D., Jiang, H., Shen, J., Melcher, K., Chen, H., Jiang, Y., Cheng, X., Wang, M.-W., Xie, X., Xu, H.E., 2022. Molecular recognition of morphine and fentanyl by the human μ-opioid receptor. Cell 185, 4361–4375.e19. 10.1016/j.cell.2022.09.041

Zivanov, J., Nakane, T., Forsberg, B.O., Kimanius, D., Hagen, W.J., Lindahl, E., Scheres, S.H., 2018. New tools for automated high-resolution cryo-EM structure determination in RELION-3. eLife 7, e42166. 10.7554/eLife.42166

