## Supplementary items for "Structural insights into the agonist selectivity and structure-based engineering of the adenosine A3 receptor"

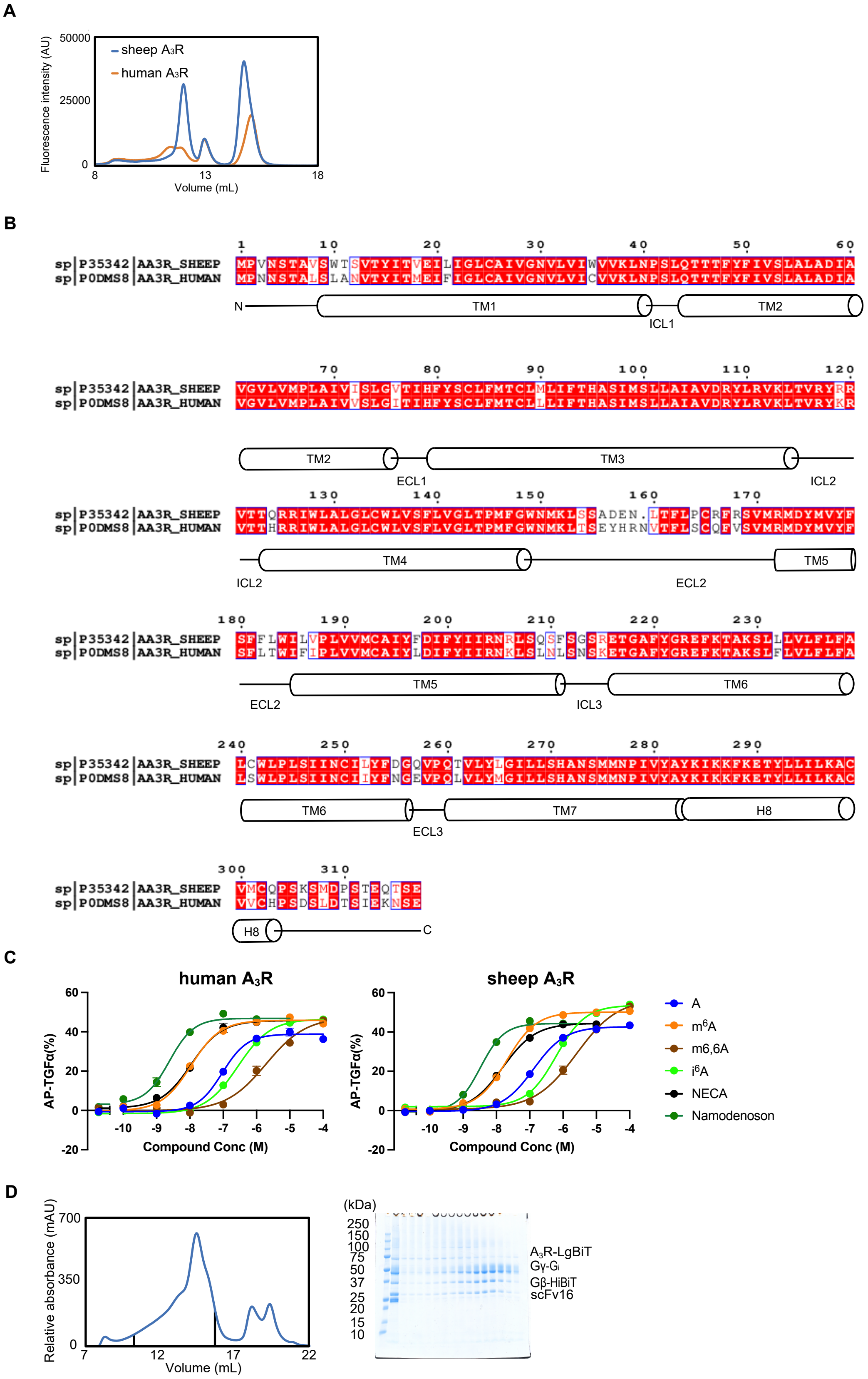


**Figure S1. Comparison of $A_3$Rs and sample preparation**

(A) Fluorescence-detection size-exclusion chromatography (FSEC) analysis of the complex formations by sheep and human $A_3$R.

(B) Alignment of the sheep and human $A_3$R amino acid sequences. Secondary structures are represented by schematic models (based on GPCRdb).

(C) TGF-α shedding assays of sheep and human $A_3$R. Both $A_3$Rs exhibited similar ligand binding profiles.

(D) Size-exclusion chromatography elution profiles of $A_3$R and SDS-PAGE analysis of the gel filtration fractions. Corresponding fractions are represented by black lines.

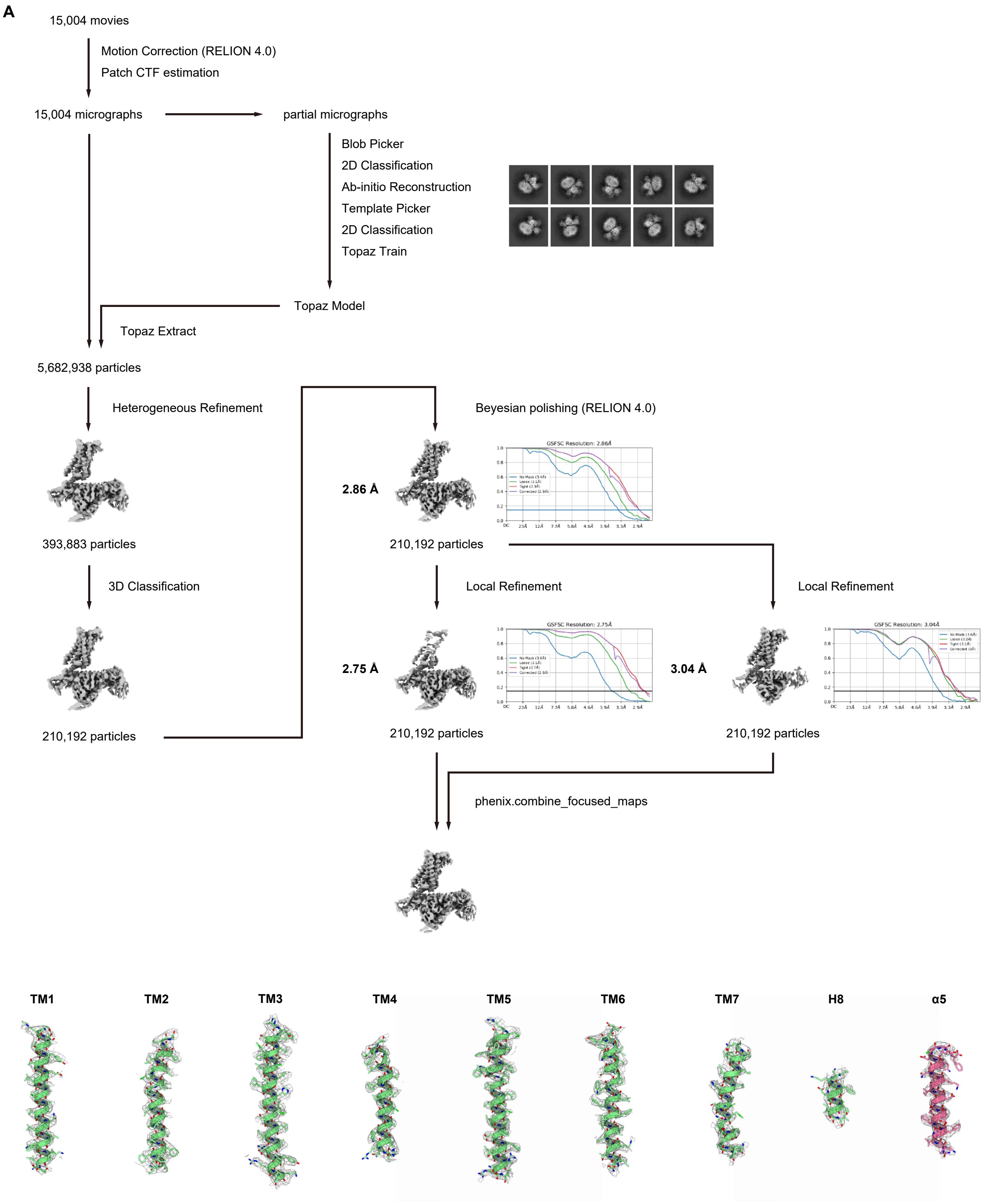
A
15,004 movies
Motion Correction (RELION 4.0)
Patch CTF estimation
15,004 micrographs
partial micrographs
Blob Picker
2D Classification
Ab-initio Reconstruction
Template Picker
2D Classification
Topaz Train
Topaz Model
Topaz Extract
5,682,938 particles
Heterogeneous Refinement
393,883 particles
3D Classification
210,192 particles
Beyesian polishing (RELION 4.0)
2.86 Å
GSFSC Resolution: 2.86Å
210,192 particles
Local Refinement
Local Refinement
2.75 Å
GSFSC Resolution: 2.75Å
3.04 Å
GSFSC Resolution: 3.04Å
210,192 particles
210,192 particles
phenix.combine_focused_maps
TM1
TM2
TM3
TM4
TM5
TM6
TM7
H8
α5

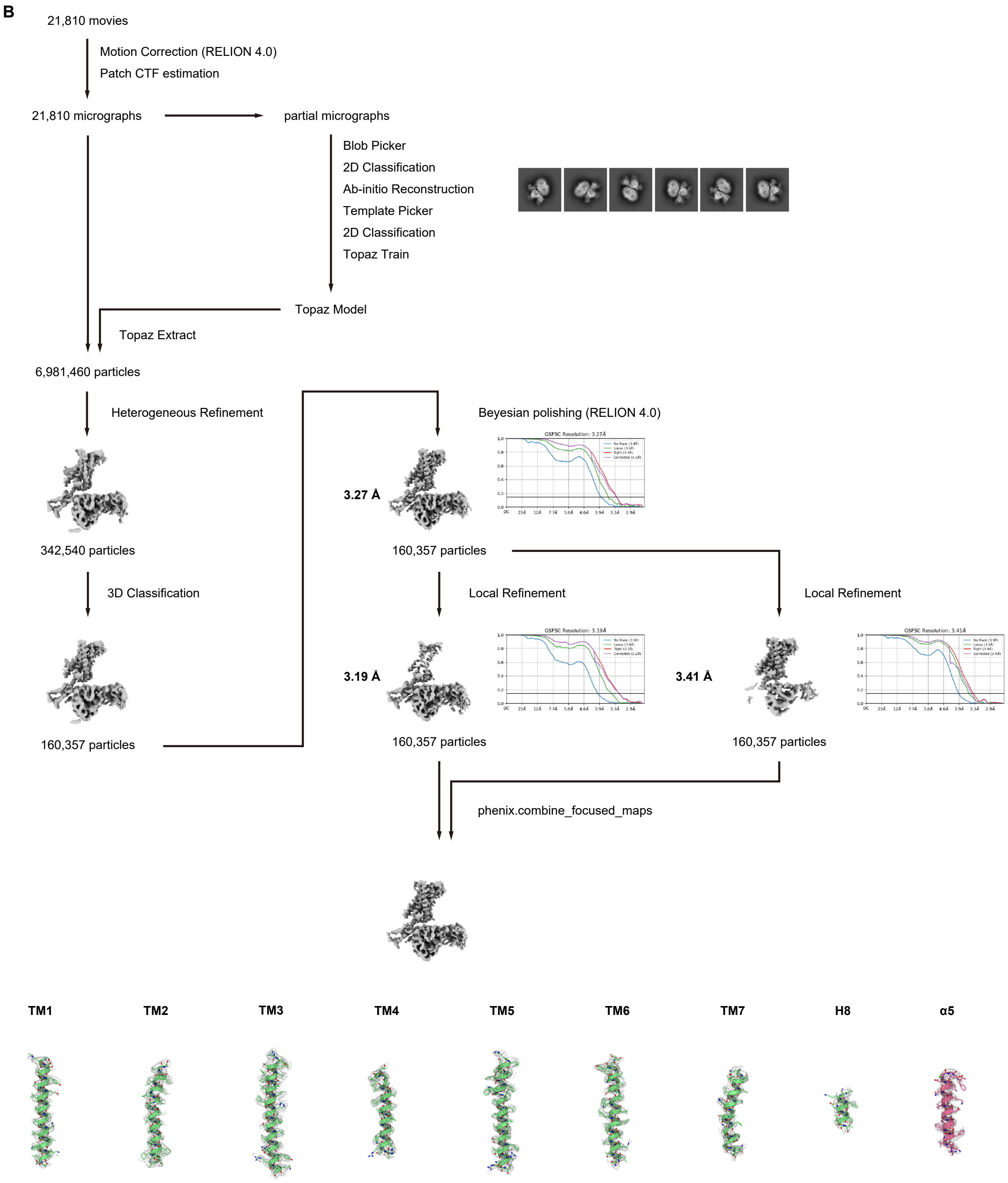
B
21,810 movies
Motion Correction (RELION 4.0)
Patch CTF estimation
21,810 micrographs
partial micrographs
Blob Picker
2D Classification
Ab-initio Reconstruction
Template Picker
2D Classification
Topaz Train
Topaz Model
Topaz Extract
6,981,460 particles
Heterogeneous Refinement
342,540 particles
3D Classification
160,357 particles
Beyesian polishing (RELION 4.0)
3.27 Å
160,357 particles
Local Refinement
Local Refinement
3.19 Å
160,357 particles
3.41 Å
160,357 particles
phenix.combine_focused_maps
TM1
TM2
TM3
TM4
TM5
TM6
TM7
H8
α5

C

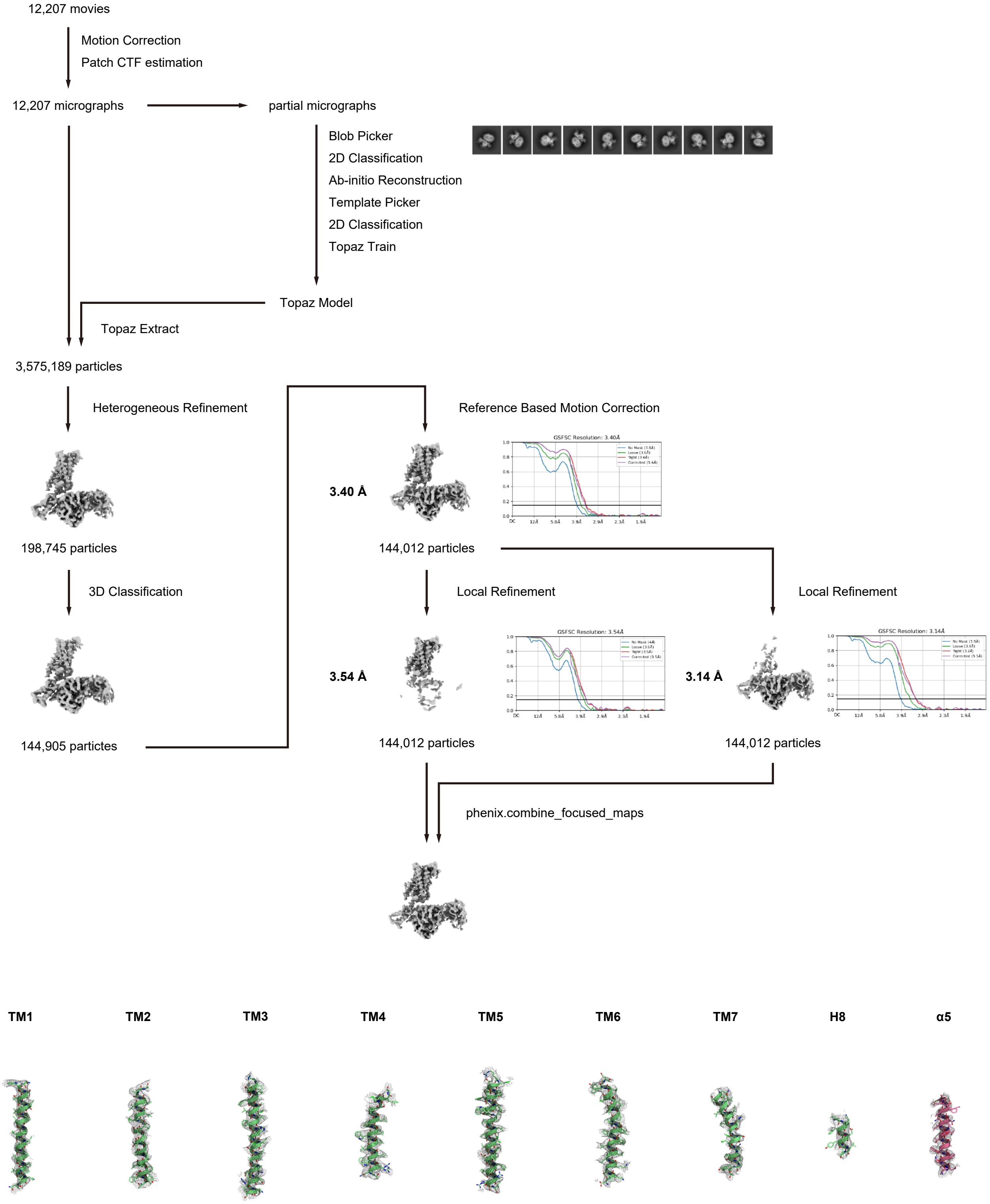

12,207 movies
Motion Correction
Patch CTF estimation
12,207 micrographs
partial micrographs
Blob Picker
2D Classification
Ab-initio Reconstruction
Template Picker
2D Classification
Topaz Train
Topaz Model
Topaz Extract
3,575,189 particles
Heterogeneous Refinement
198,745 particles
3D Classification
144,905 particles
Reference Based Motion Correction
3.40 Å
GSFSC Resolution: 3.40Å
144,012 particles
Local Refinement
Local Refinement
3.54 Å
GSFSC Resolution: 3.54Å
3.14 Å
GSFSC Resolution: 3.14Å
144,012 particles
144,012 particles
phenix.combine_focused_maps
TM1
TM2
TM3
TM4
TM5
TM6
TM7
H8
α5

D

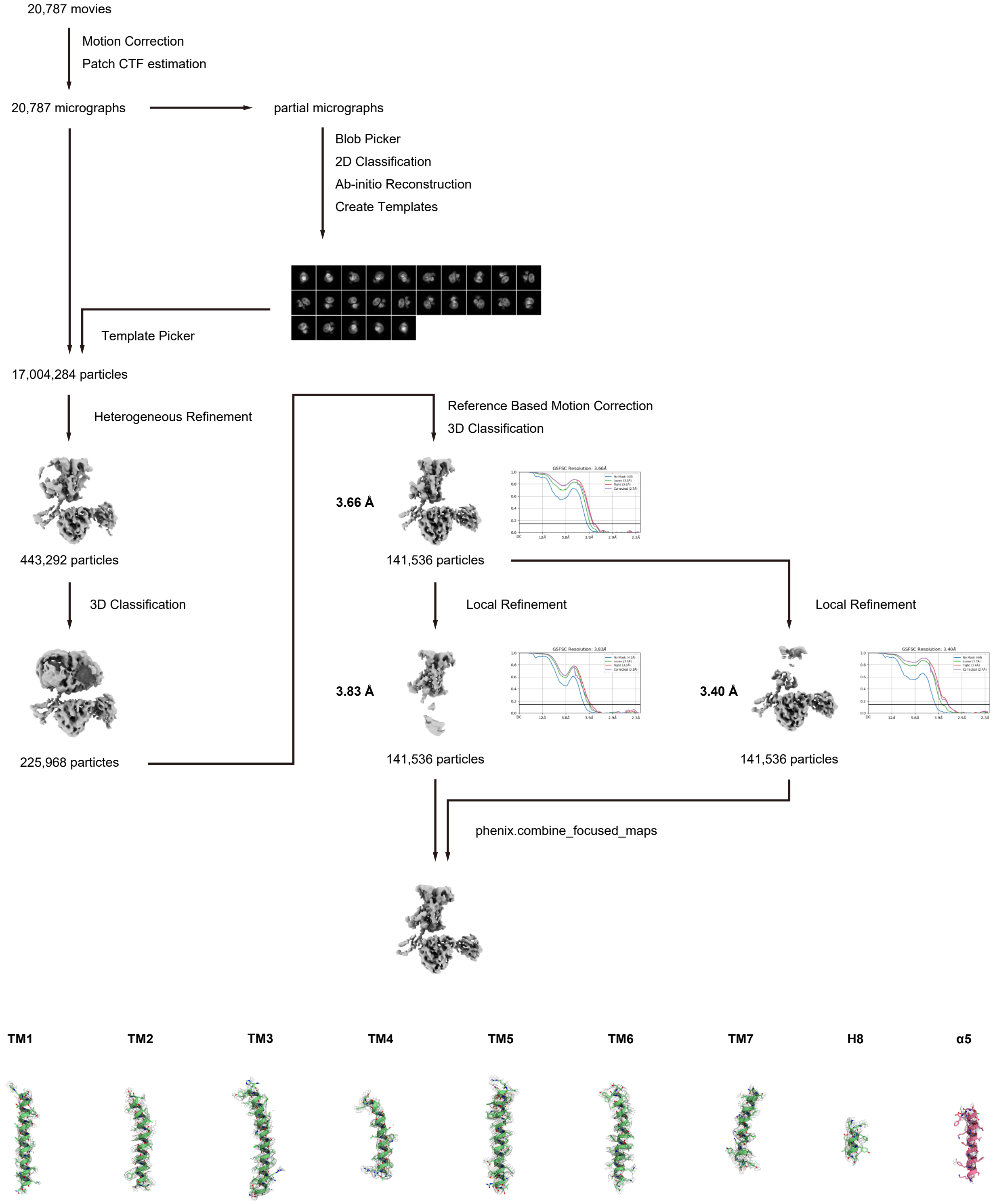
20,787 movies
Motion Correction
Patch CTF estimation
20,787 micrographs
partial micrographs
Blob Picker
2D Classification
Ab-initio Reconstruction
Create Templates
Template Picker
17,004,284 particles
Heterogeneous Refinement
443,292 particles
3D Classification
225,968 particles
Reference Based Motion Correction
3D Classification
3.66 Å
141,536 particles
Local Refinement
3.83 Å
141,536 particles
Local Refinement
3.40 Å
141,536 particles
phenix.combine_focused_maps
TM1
TM2
TM3
TM4
TM5
TM6
TM7
H8
α5

**E**

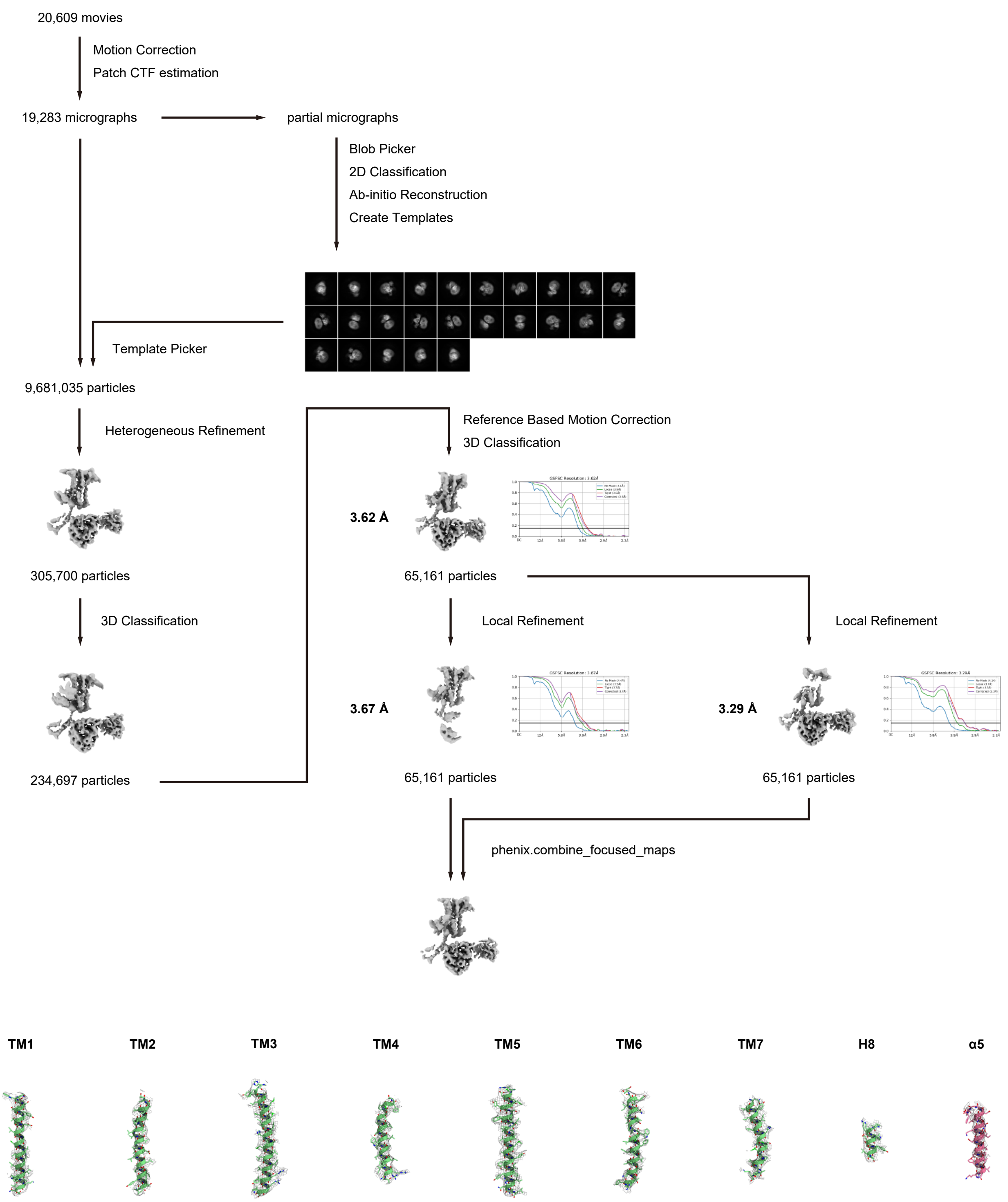


**Figure S2. Cryo-EM workflow, maps, and model quality**

(A-E) Flow charts of the cryo-EM data processing for the $A_3R$-$G_i$ complexes bound to NECA (A), adenosine (B), $m^6A$ (C), $i^6A$ (D), and namodenoson (E). Unless otherwise noted, analyses were performed with cryoSPARC.

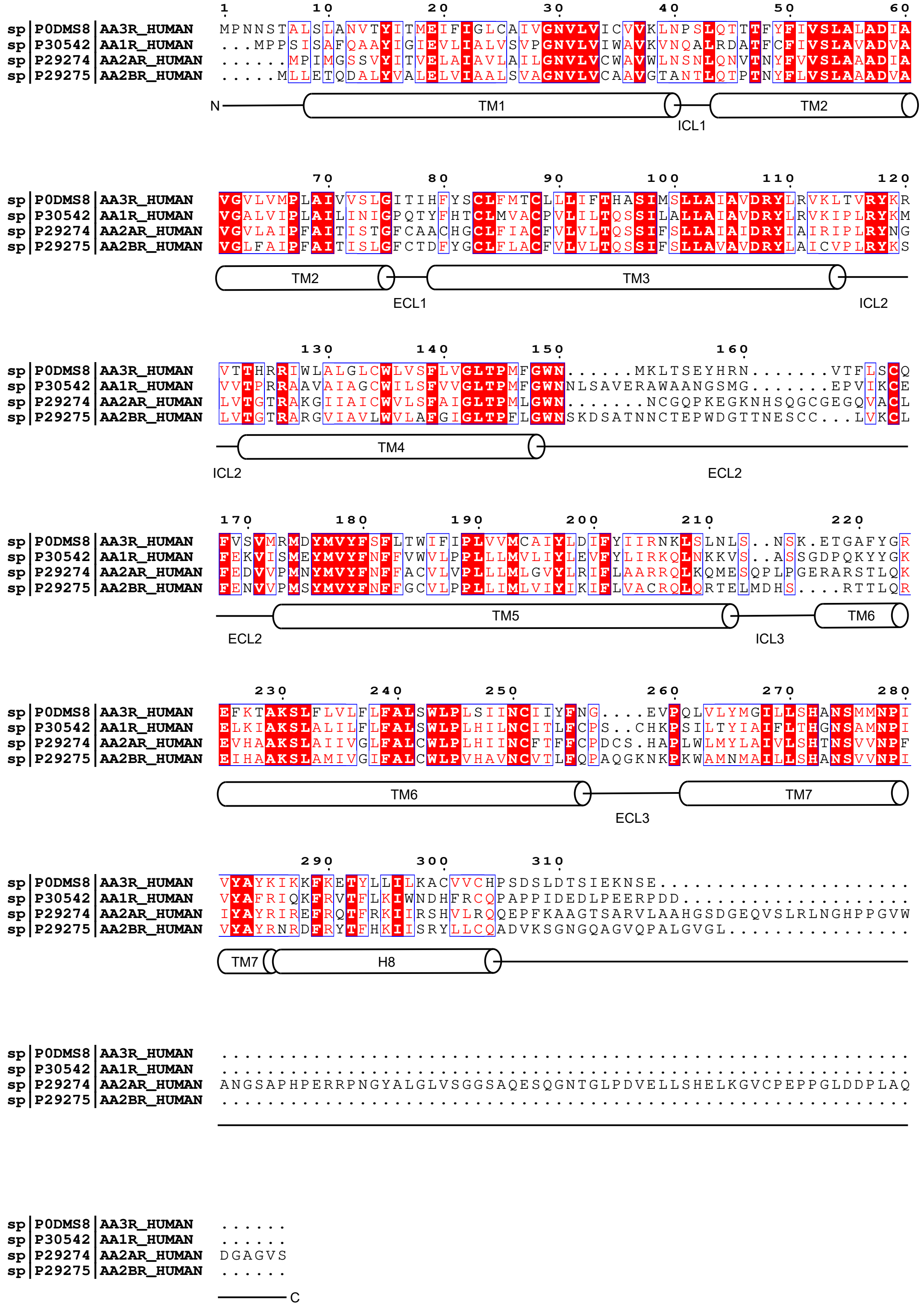


**Figure S3. Alignment of the adenosine receptors**

Amino acid alignment of the adenosine receptors. Secondary structures are represented by schematic models (based on GPCRdb).

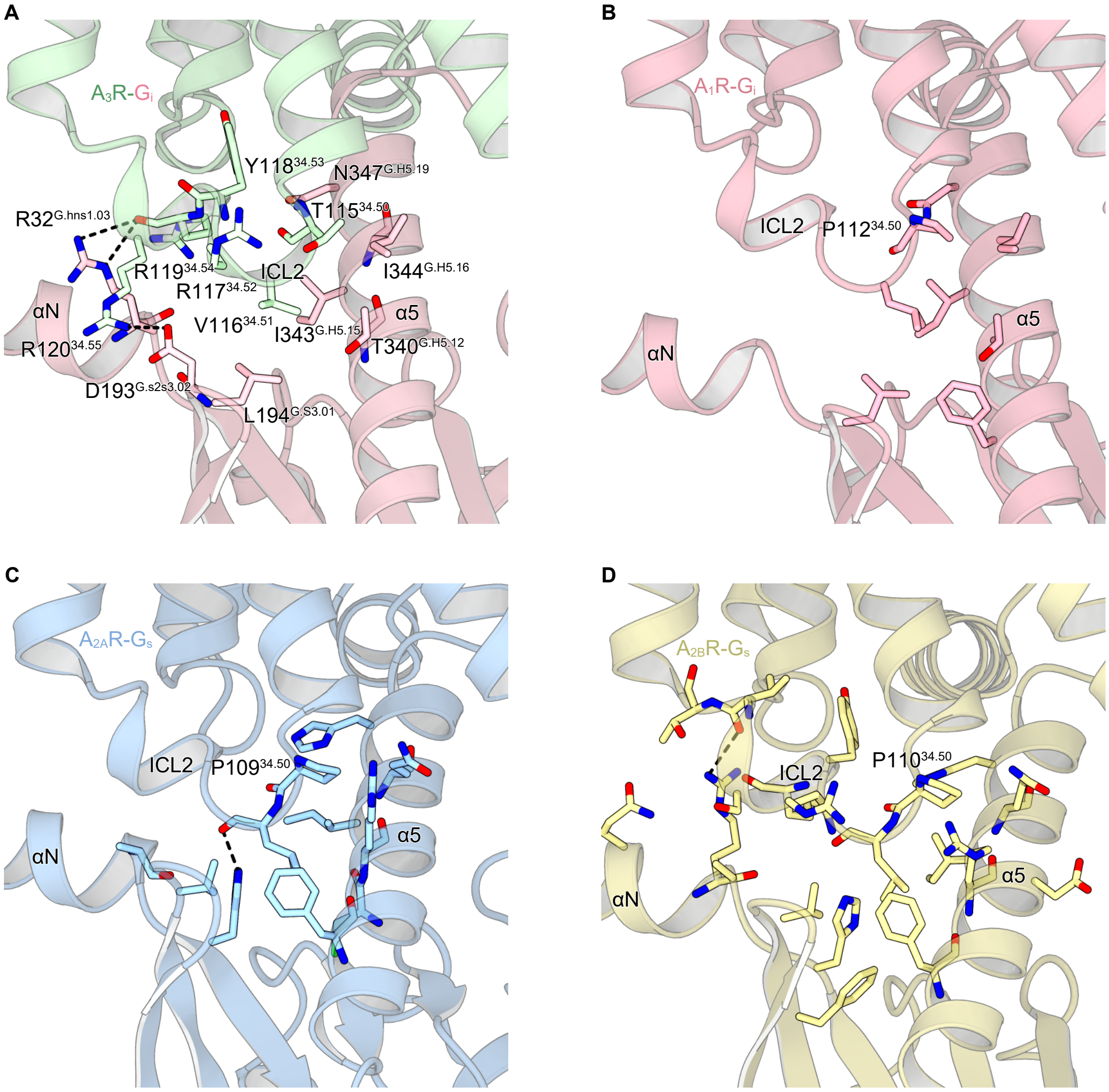


**Figure S4. Comparison of ECL2s of adenosine receptors**

(A-D) Receptor-G-protein interactions of $A_1R$ (A; PDB 6D9H), $A_{2A}R$ (B; PDB 6GDG), $A_{2B}R$ (C; PDB 7XY7), and $A_3R$ (D). $A_3R$ forms extensive interactions with $G_i$. This may be a consequence of $T115^{34.50}$, which endows $A_3R$ with a slightly elongated helix compared to the other adenosine receptors with $P^{34.50}$.

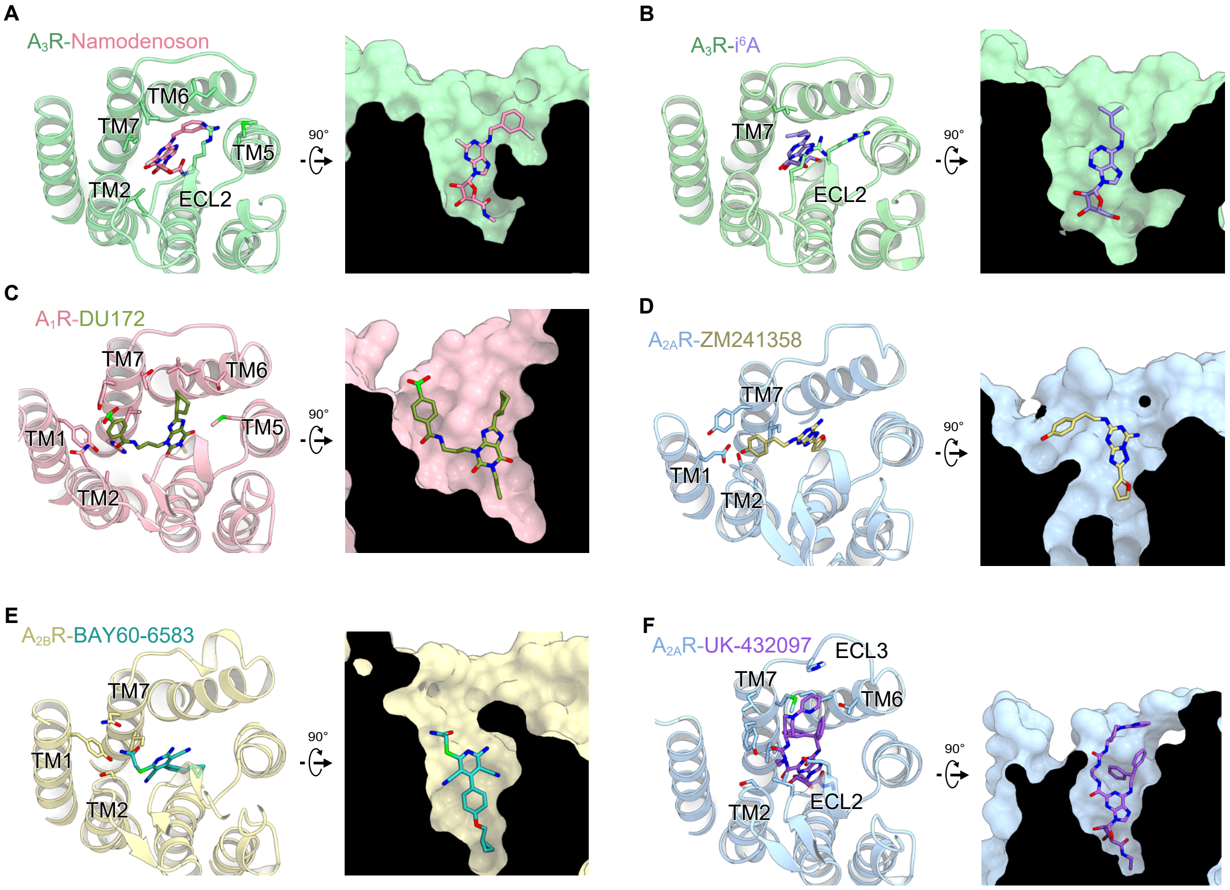


**Figure S5. Binding modes of selective ligands**

(A-F) Binding modes of the adenosine receptors and their selective ligands (left), and cross sections of the orthosteric pockets (right). (A) Namodenoson-bound $A_3R$. (B) $i^6A$-bound $A_3R$. (C) DU172-bound $A_1R$ (PDB 5UEN). (D) ZM241358-bound $A_{2A}R$ (PDB 3PWH). (E) BAY60-6583-bound $A_{2B}R$ (PDB 7XY6). (F) UK-432097-bound $A_{2A}R$ (PDB 3QAK). While occupying the orthosteric pocket, every ligand exhibits interactions with the extracellular portion of the receptor. These extracellular interactions should be essential for the selective activation of an adenosine receptor.

### Table S1. Comprehensive screening of modified nucleosides against human A3R

| Relative GPCR activation of each nucleoside (TPA = 1 | t6A | ms2t6A | ms2i6A | m6Am | Am | m6A | m6,6A | i6A | m2,8A | N6-sucA | ms2A | m2A | Im | m1I | m2,2G | m2G | m1G | Gm | m7G | 8OHG | ac4C |
|---|---|---|---|---|---|---|---|---|---|---|---|---|---|---|---|---|---|---|---|---|---|
| hA1R +Gq/o Dabsorbance (nucleoside = 100nM) | 0.00 | -0.02 | 0.03 | 0.06 | 0.03 | 0.06 | 0.02 | 0.06 | -0.05 | -0.06 | -0.06 | 0.17 | -0.03 | 0.03 | -0.08 | -0.01 | 0.00 | -0.03 | -0.09 | -0.11 | 0.02 |
| hA2AR +Gq/s Dabsorbance (nucleoside = 1μM) | -0.01 | 0.01 | 0.00 | -0.02 | -0.01 | 0.05 | 0.10 | 0.05 | -0.01 | 0.04 | 0.02 | 0.15 | 0.03 | -0.01 | 0.00 | 0.03 | 0.01 | 0.03 | 0.01 | -0.01 | 0.01 |
| hA2BR +Gq/s Dabsorbance (nucleoside = 5μM) | 0.01 | 0.02 | 0.00 | 0.01 | 0.02 | 0.13 | 0.07 | 0.14 | 0.01 | 0.01 | -0.01 | 0.24 | 0.02 | -0.01 | 0.01 | -0.01 | -0.01 | -0.01 | 0.01 | 0.00 | 0.03 |
| hA3R +Gq/i3 Dabsorbance (nucleoside = 100nM) | -0.04 | -0.05 | -0.04 | 0.04 | -0.01 | 0.92 | -0.02 | 0.19 | -0.03 | -0.06 | -0.01 | 0.34 | -0.05 | -0.03 | -0.01 | -0.01 | -0.01 | 0.03 | -0.01 | -0.02 | -0.03 |

| Relative GPCR activation of each nucleoside (TPA = 1 | Cm | m5C | f5C | m3C | m5Cm | hm5C | s2C | m4Cm | Y | acp3U | D | Um | m3U | m5U | tm5U | s2U | tm5s2U | m5Um | m1Y | s4U | ho5U |
|---|---|---|---|---|---|---|---|---|---|---|---|---|---|---|---|---|---|---|---|---|---|
| hA1R +Gq/o Dabsorbance (nucleoside = 100nM) | -0.04 | -0.06 | -0.05 | -0.04 | -0.04 | -0.02 | -0.01 | -0.07 | -0.04 | -0.07 | -0.08 | -0.09 | -0.06 | -0.05 | -0.06 | -0.08 | -0.06 | 0.00 | 0.00 | -0.03 | -0.01 |
| hA2AR +Gq/s Dabsorbance (nucleoside = 1μM) | 0.01 | 0.02 | 0.02 | 0.02 | 0.00 | 0.02 | 0.05 | 0.05 | 0.03 | 0.01 | -0.03 | -0.01 | 0.00 | 0.00 | 0.07 | 0.01 | -0.02 | 0.01 | 0.03 | 0.01 | 0.00 |
| hA2BR +Gq/s Dabsorbance (nucleoside = 5μM) | 0.01 | 0.00 | -0.01 | 0.01 | 0.00 | 0.02 | -0.01 | -0.02 | 0.00 | 0.00 | 0.01 | 0.01 | -0.01 | 0.01 | -0.01 | 0.01 | 0.02 | 0.01 | -0.01 | 0.01 | 0.01 |
| hA3R +Gq/i3 Dabsorbance (nucleoside = 100nM) | -0.04 | -0.02 | -0.01 | -0.04 | -0.01 | -0.01 | -0.06 | -0.05 | 0.00 | 0.01 | -0.03 | -0.07 | -0.05 | -0.04 | -0.08 | -0.08 | -0.05 | 0.00 | -0.02 | -0.05 | -0.04 |

A3R activation (%AP-TGFα release)

| compound concentration Log10 (M) | Adenosine | | | | | | m6A | | | | | | m6,6A | | | i6A | | | m6A (MOCK) | | | m6,6A(MOCK) | | | i6A(MOCK) | | | t6A(MOCK) | | |
|---|---|---|---|---|---|---|---|---|---|---|---|---|---|---|---|---|---|---|---|---|---|---|---|---|---|---|---|---|---|---|
| -13 | 2.17 | -0.26 | 0.18 | -1.63 | -1.5 | -2.78 | -2.83 | 1.01 | -1.16 | 0.46 | -0.51 | 0.59 | 0.91 | 1.91 | 1.2 | 0.6 | -1 | -2.72 | 0.64 | -0.53 | -0.12 | 0.59 | 0.2 | -0.79 | 0.19 | 0.25 | -0.44 | 0 | 0 | 0 |
| -10 | | | | 3.13 | -1.1 | 1.2 | | | | 3.54 | -0.5 | 2.78 | | | | | | | | | | | | | | | | | | |
| -9 | 1.43 | -5.91 | -3.89 | -1.57 | -0.4 | 0.7 | 4.75 | -2.56 | 3.45 | 5.7 | -0.49 | 5.84 | -1.96 | 3.13 | 2.24 | -2.98 | 1.17 | -1.69 | -0.67 | -0.18 | 0.23 | -1.69 | 0.7 | -0.81 | 0.58 | -0.27 | -0.92 | 0 | 0 | 0 |
| -8 | 1.69 | 2.74 | -0.66 | 2.05 | 5.3 | 3.7 | 18.5 | 18.3 | 24.8 | 25.2 | 22.7 | 25.5 | -0.38 | -1.47 | -1.66 | -2.21 | -1.19 | 2.27 | -0.25 | 0.27 | 0.5 | -0.05 | -0.7 | -0.11 | 0.32 | -0.16 | 0.06 | 0 | 0 | 0 |
| -7 | 21.8 | 22 | 14.7 | 20.1 | 16.1 | 23.6 | 36.9 | 40.2 | 36.3 | 44.3 | 41.3 | 44 | 4.55 | 0.79 | 3.5 | 15.4 | 11.9 | 11 | 0.22 | 0.5 | 0.06 | 0.2 | 1.46 | 0.87 | 0.37 | 0.9 | -0.56 | 0 | 0 | 0 |
| -6 | 32.1 | 32.3 | 32.7 | 41.2 | 38.5 | 42.7 | 42 | 38.9 | 48.1 | 50.4 | 44.7 | 46 | 21.2 | 15.6 | 23.6 | 36.4 | 36.6 | 30.6 | 1.53 | 2.03 | 0.79 | 5.52 | 4.3 | 5.62 | 1.76 | 2.29 | 0.06 | 2.63536 | 4.18487 | 2.1591 |
| -5 | 35.5 | 34.6 | 35.9 | 44.6 | 44.9 | 43.8 | 43.3 | 42.2 | 47.2 | 53.9 | 46.4 | 51.2 | 33.4 | 34.8 | 35.2 | 45 | 46.6 | 42.6 | 2.26 | 2.41 | 2.75 | 12.4 | 11.7 | 9.29 | 6.97 | 6.16 | 6.3 | 0.19127 | 0.91968 | -0.7936 |
| -4 | 38.7 | 33.8 | 36.4 | | | | 46.6 | 40.4 | 45.5 | | | | 44.4 | 45.6 | 47.4 | 49 | 44.4 | 45.4 | 4.09 | 3.76 | 3.91 | 20.3 | 20.2 | 21.3 | 13 | 12.3 | 10.8 | 0 | 0 | 0 |

### Table S2. Analyses human and sheep $A_3$Rs using the TGFα shedding assay

| | A | | | | m6A | | | | m6,6A | | | | i6A | | | | NECA | | | | Namodenoson | | | |
|---|---|---|---|---|---|---|---|---|---|---|---|---|---|---|---|---|---|---|---|---|---|---|---|---|
| | pEC50 | | Emax | | pEC50 | | Emax | | pEC50 | | Emax | | pEC50 | | Emax | | pEC50 | | Emax | | pEC50 | | Emax | |
| | Mean ± SEM | EC50 (nM) | Mean ± SEM | n | Mean ± SEM | EC50 (nM) | Mean ± SEM | n | Mean ± SEM | EC50 (nM) | Mean ± SEM | n | Mean ± SEM | EC50 (nM) | Mean ± SEM | n | Mean ± SEM | EC50 (nM) | Mean ± SEM | n | Mean ± SEM | EC50 (nM) | Mean ± SEM | n |
| Human A3R | 7.0 ± 0.079 | 110 | 41 ± 2.2 | 6 | 8.0 ± 0.023 | 11 | 47 ± 1.5 | 6 | 5.7 ± 0.085 | 1900 | 47 ± 0.53 | 3 | 6.6 ± 0.062 | 290 | 48 ± 1.1 | 3 | 7.9 ± 0.040 | 11 | 44 ± 0.62 | 3 | 8.6 ± 0.057 | 2.5 | 44 ± 0.60 | 3 |
| Sheep A3R | 6.9 ± 0.043 | 130 | 43 ± 1.0 | 6 | 6.9 ± 0.043 | 20 | 49 ± 0.71 | 6 | 5.6± 0.090 | 2600 | 56 ± 3.0 | 3 | 6.2 ± 0.061 | 640 | 52 ± 2.1 | 3 | 7.8 ± 0.015 | 16 | 45 ± 0.63 | 3 | 8.5 ± 0.064 | 3.4 | 45 ± 0.58 | 3 |

### Table S3. Mutational analyses using the TGFα shedding assay

| | A | | | | m6A | | | | m6,6A | | | | i6A | | | |
|---|---|---|---|---|---|---|---|---|---|---|---|---|---|---|---|---|
| | pEC50 | | Emax | | pEC50 | | Emax | | pEC50 | | Emax | | pEC50 | | Emax | |
| | Mean ± SEM | EC50 (nM) | Mean ± SEM | n | Mean ± SEM | EC50 (nM) | Mean ± SEM | n | Mean ± SEM | EC50 (nM) | Mean ± SEM | n | Mean ± SEM | EC50 (nM) | Mean ± SEM | n |
| A3R-WT | 6.9 ± 0.039 | 130 | 43 ± 1.0 | 12 | 7.9 ± 0.034 | 14 | 49 ± 1.0 | 12 | 5.4 ± 0.022 | 3900 | 57 ± 0.65 | 12 | 6.3 ± 0.025 | 480 | 52 ± 0.60 | 16 |
| V169A | 7.0 ± 0.032 | 95 | 40 ± 0.82 | 6 | 7.7 ± 0.026 | 18 | 49 ± 0.95 | 5 | 5.6 ± 0.018 | 2400 | 55 ± 4.2 | 3 | 7.0 ± 0.071 | 98 | 52 ± 3.0 | 3 |
| M174A | 5.8 ± 0.052 | 1600 | 59 ± 1.6 | 3 | 6.6 ± 0.023 | 280 | 64 ± 1.7 | 3 | NA | NA | NA | 3 | 7.4 ± 0.017 | 36 | 70 ± 4.5 | 3 |
| M174L | 6.6 ± 0.061 | 260 | 53 ± 1.6 | 3 | 7.8 ± 0.11 | 16 | 59 ± 1.1 | 3 | 5.3 ± 0.018 | 5400 | 62 ± 1.8 | 3 | 6.2 ± 0.018 | 660 | 60 ± 3.1 | 3 |
| I253A | 6.5 ± 0.050 | 320 | 51 ± 1.2 | 3 | 7.0 ± 0.064 | 110 | 58 ± 1.3 | 3 | 4.8 ± 0.021 | 17000 | 70 ± 0.61 | 3 | 7.4 ± 0.063 | 41 | 64 ± 1.2 | 3 |
| L264A | 6.7 ± 0.11 | 220 | 50 ± 0.71 | 4 | 7.9 ± 0.078 | 12 | 57 ± 1.4 | 3 | 5.4 ± 0.040 | 4000 | 65 ± 2.3 | 3 | 7.2 ± 0.044 | 69 | 63 ± 1.2 | 3 |
| V169L | 7.0 ± 0.023 | 97 | 41 ± 0.66 | 9 | 7.7 ± 0.037 | 18 | 49 ± 1.6 | 6 | 5.6 ± 0.017 | 2300 | 52 ± 2.4 | 3 | 5.9 ± 0.027 | 1200 | 54 ± 3.9 | 3 |
| V169F | 6.7 ± 0.053 | 210 | 53 ± 1.7 | 3 | 7.0 ± 0.045 | 100 | 59 ± 2.6 | 3 | 4.5 ± 0.16 | 35000 | 80 ± 5.9 | 3 | 5.9 ± 0.044 | 1300 | 60 ± 1.8 | 5 |
| M174F | 6.5 ± 0.057 | 360 | 56 ± 2.4 | 3 | 7.1 ± 0.046 | 74 | 60 ± 0.66 | 3 | 4.5 ± 0.0092 | 35000 | 82 ± 1.7 | 3 | 5.4 ± 0.029 | 4200 | 64 ± 1.5 | 3 |
| I253F | 6.5 ± 0.16 | 330 | 51 ± 0.95 | 3 | 7.4 ± 0.18 | 51 | 56 ± 2.2 | 3 | 4.9 ± 0.11 | 13000 | 62 ± 0.57 | 3 | 7.1 ± 0.18 | 85 | 62 ± 1.6 | 3 |
| I253M | 6.4 ± 0.038 | 400 | 57 ± 1.5 | 3 | 7.3 ± 0.037 | 47 | 62 ± 1.6 | 4 | 4.8 ± 0.047 | 16000 | 72 ± 0.12 | 3 | 6.4 ± 0.024 | 430 | 66 ± 1.9 | 3 |
| L264F | 5.1 ± 0.064 | 8600 | 56 ± 1.6 | 3 | 6.0 ± 0.019 | 1100 | 64 ± 1.9 | 3 | NA | NA | NA | 3 | 5.5 ± 0.024 | 3400 | 70 ± 1.0 | 3 |
| L264M | 6.8 ± 0.042 | 180 | 43 ± 0.75 | 4 | 7.3 ± 0.060 | 48 | 51 ± 0.76 | 3 | 4.9 ± 0.034 | 11000 | 63 ± 2.2 | 3 | 6.0 ± 0.0070 | 1100 | 58 ± 1.2 | 3 |
| V169T | 7.0 ± 0.030 | 94 | 40 ± 1.1 | 5 | 8.0 ± 0.046 | 11 | 43 ± 1.1 | 4 | 5.6 ± 0.023 | 2400 | 52 ± 0.93 | 3 | 6.3 ± 0.047 | 470 | 50 ± 1.8 | 3 |
| M174Y | 5.4 ± 0.17 | 4400 | 65 ± 1.8 | 3 | 5.7 ± 0.015 | 1800 | 64 ± 2.5 | 3 | 5.1 ± 0.050 | 8500 | 41 ± 10 | 4 | 5.2 ± 0.039 | 6200 | 67 ± 3.0 | 3 |
| I253N | 6.8 ± 0.041 | 170 | 51 ± 1.2 | 5 | 7.2 ± 0.042 | 69 | 55 ± 2.4 | 4 | 4.8 ± 0.11 | 15000 | 70 ± 1.6 | 3 | 7.3 ± 0.040 | 45 | 60 ± 1.5 | 3 |
| L264N | 6.1 ± 0.081 | 790 | 58 ± 0.33 | 3 | 7.1 ± 0.051 | 81 | 62 ± 2.1 | 3 | 4.2 ± 0.26 | 98000 | 100 ± 14 | 3 | 5.8 ± 0.0043 | 1600 | 66 ± 0.91 | 3 |
| V169N | 6.7 ± 0.041 | 190 | 54 ± 0.61 | 4 | 7.2 ± 0.17 | 75 | 55 ± 1.1 | 4 | 5.2 ± 0.014 | 5600 | 64 ± 0.33 | 3 | 6.2 ± 0.28 | 1400 | 60 ± 1.1 | 4 |
| I253Q | 6.3 ± 0.058 | 500 | 57 ± 1.3 | 3 | 7.0 ± 0.085 | 99 | 62 ± 1.4 | 3 | 4.4 ± 0.11 | 46000 | 89 ± 8.7 | 3 | 7.1 ± 0.015 | 80 | 68 ± 2.3 | 3 |
| L264Q | 6.7 ± 0.061 | 230 | 52 ± 0.53 | 3 | 8.0 ± 0.044 | 9.5 | 57 ± 2.1 | 3 | 5.5 ± 0.069 | 3500 | 67 ± 3.1 | 3 | 6.6± 0.051 | 267 | 63 ± 1.4 | 4 |
| I253N+V169N | 6.9 ± 0.033 | 130 | 47 ± 0.54 | 3 | 6.8 ± 0.032 | 170 | 53 ± 0.57 | 3 | 4.8 ± 0.24 | 19000 | 58 ± 6.3 | 3 | 6.9 ± 0.075 | 140 | 59 ± 1.0 | 3 |
| I253N+V169F | 6.3 ± 0.011 | 470 | 51 ± 0.86 | 3 | 6.2 ± 0.018 | 680 | 55 ± 1.9 | 3 | 4.6 ± 0.29 | 36000 | 44 ± 9.2 | 3 | 6.1 ± 0.015 | 830 | 62 ± 1.3 | 3 |
| I253Q+V169N | 5.5 ± 0.054 | 3200 | 13 ± 0.48 | 3 | 5.4 ± 0.034 | 4400 | 17 ± 1.9 | 3 | 4.6 ± 0.13 | 26000 | 29 ± 3.1 | 3 | 5.1 ± 0.061 | 8400 | 31 ± 1.6 | 3 |
| I253Q+V169F | 6.1 ± 0.031 | 720 | 45 ± 0.46 | 3 | 6.0 ± 0.038 | 1100 | 53 ± 1.9 | 3 | 4.7 ± 0.082 | 21000 | 33 ± 0.34 | 3 | 5.7 ± 0.032 | 1800 | 61 ± 0.26 | 3 |

### Table S4. Details of each tested nucleoside

| compounds | abbreviation | source | identifier |
|---|---|---|---|
| $N^{6}$-Threonylcarbamoyladenosine | t6A | Wako | custom-synthesized |
| 2-Methylthio-$N^{6}$-threonylcarbamoyladenosine | ms2t6A | Wako | custom-synthesized; CAS: 70333-82-3 |
| 2-Methylthio-$N^{6}$-isopentenyladenosine | ms2i6A | Wako | custom-synthesized; CAS: 20859-00-1 |
| $N^{6}$, 2'-*O*-Dimethyladenosine | m6Am | Toronto Research Chemicals | Cat#D447415; CAS:57817-83-1 |
| 2'-*O*-Methyladenosine | Am | Tokyo Chemical Industry | Cat#M2291; CAS:2140-79-6 |
| 2'-*O*-Methylinosine | Im | Santa Cruz Biotechnology | Cat#sc-283498; CAS:3881-21-8 |
| 1-Methylinosine | m1I | Santa Cruz Biotechnology | Cat#sc-483758; CAS:2140-73-0 |
| $N^{6}$-Methyladenosine | m6A | Abcam | Cat#ab145715; CAS:1867-73-8 |
| $N^{6}$,$N^{6}$-Dimethyladenosine | m6,6A | MedChem Express | Cat#HY-101984; CAS:2620-62-4 |
| $N^{6}$-Isopentenyladenosine | i6A | Wako | custom-synthesized |
| $N^{4}$-Acetylcytidine | ac4C | Combi-Blocks | Cat#QB-9019; CAS:3768-18-1 |
| 2'-*O*-Methylcytidine | Cm | Tokyo Chemical Industry | Cat#M2317; CAS:2140-72-9 |
| 5-Methylcytidine | m5C | Tokyo Chemical Industry | Cat#M1931; CAS:2140-61-6 |
| 5-Formylcytidine | f5C | Toronto Research Chemicals | Cat#F698965; CAS:148608-53-1 |
| 3-Methylcytidine | m3C | MedChem Express | Cat#HY-111645; CAS: 2140-64-9 |
| 5,2'-O-Dimethylcytidine | m5Cm | Glanlen | Cat#GL100726; CAS: 113886-70-7 |
| 5-Hydroxymethylcytidine | hm5C | Tokyo Chemical Industry | Cat#H1863; CAS: 19235-17-7 |
| $N^{2}$,$N^{2}$-Dimethylguanosine | m2,2G | MedChem Express | Cat#HY-113137; CAS:2140-67-2 |
| $N^{2}$-Methylguanosine | m2G | Biosynth Carbosynth | Cat#NM35522; CAS:2140-77-4 |
| 1-Methylguanosine | m1G | Santa Cruz Biotechnology | Cat#sc-500889; CAS:2140-65-0 |
| 2'-*O*-Methylguanosine | Gm | Tokyo Chemical Industry | Cat#M2318; CAS:2140-71-8 |
| 7-Methylguanosine | m7G | Sigma-Aldrich | Cat#M0627; CAS:20244-86-4 |
| Pseudouridine | Ψ | Tokyo Chemical Industry | Cat#P2396; CAS:1445-07-4 |
| 3-(3-Amino-3-carboxypropyl) uridine | acp3U | Alsachim | custom-synthesized |
| 5,6-Dihydrouridine | D | Toronto Research Chemicals | Cat#D449668; CAS:5627-05-4 |
| 2'-*O*-Methyluridine | Um | Tokyo Chemical Industry | Cat#M2290; CAS:2140-76-3 |
| 3-Methyluridine | m3U | Sigma-Aldrich | Cat#M4129; CAS:2140-69-4 |
| 5−Methyluridine | m5U | Tokyo Chemical Industry | Cat#M1405; CAS:1463-10-1 |
| 5-Taurinomethyluridine | τm5U | Wako | custom-synthesized; CAS: 497258-53-4 |
| 2-Thiouridine | s2U | Tokyo Chemical Industry | Cat#T4108; CAS: 20235-78-3 |
| 5-Taurinomethyl-2-thiouridine | τm5s2U | Wako | custom-synthesized; CAS: 497258-54-5 |
| 5,2′−O−Dimethyluridine | m5Um | Combi-Blocks | Cat#QC-7341; CAS: 55486-09-4 |
| N1-Methylpseudouridine | m1Ψ | MedChem Express | Cat#HY-112582 ; CAS: 13860-38-3 |
| 4-Thiouridine | s4U | Tokyo Chemical Industry | Cat#T4059; CAS:13957-31-8 |
| 5-Hydroxyuridine | ho5U | Tokyo Chemical Industry | Cat#H1865; CAS: 957-77-7 |
| 2-Methyladenosine | m2A | Santa Cruz Biotechnology | Cat#sc-500888; CAS: 16526-56-0 |
| 8-Hydroxyguanosine | 8OHG | Cayman Chemical | Cat#89300; CAS: 3868-31-3 |
| 2,8-Dimethyladenosine | m2,8A | MedChem Express | Cat#HY-152702; CAS: 63954-66-5 |
| N6-Succinyl adenosine | N6-sucA | Biosynth | Cat#NS16562; CAS: 4542-23-8 |
| 2-Methylathioadenosine | ms2A | Carbosynth | Cat#NM09610; CAS: 4105-39-9 |
| 2-Thiocytidine | s2C | Santa Cruz Biotechnology | Cat#sc-283290; CAS: 13239-97-9 |
| N4,2′−O−dimethylcytidine | m4Cm | Biosynth | Cat#NM144592; CAS: 13048-95-8 |
